# Skin microbiomes of frogs vary among individuals and body regions, revealing differences that reflect known patterns of chytrid infection

**DOI:** 10.1101/2025.02.05.636728

**Authors:** Sonia L. Ghose, Jonathan A. Eisen

## Abstract

The amphibian skin microbiome is an important line of defense against pathogens including the deadly chytrid fungus, *Batrachochytrium dendrobatidis* (*Bd*). Intra-species variation in disease susceptibility and intra-individual variation in infection distribution across the skin, therefore, may relate to differences in skin microbiomes. However, characterization of microbiome variation within and among amphibian individuals is needed. We utilized 16S rRNA gene amplicon sequencing to compare microbiomes of ten body regions from nine captive *R. sierrae* individuals and their tank environments. While frogs harbored distinct microbial communities compared to their tank environments, tank identity was associated with more variation in frog microbiomes than individual frog identity. Within individuals, we detected differences between microbiomes of body regions where *Bd* infection would be expected compared to regions that infrequently experience infection. Notably, the bacterial families Burkholderiaceae (phylum Proteobacteria) and Rubritaleaceae (phylum Verrucomicrobia) were dominant on frog skin, and the relative abundances of undescribed members of these families were important to describing differences among and within individuals. Two undescribed Burkholderiaceae taxa were found to be putatively *Bd*-inhibitory, and both showed higher relative abundance on body regions where *Bd* infection is often localized. These findings highlight the importance of considering intrapopulation and intraindividual heterogeneities, which could provide insights relevant to predicting localized interactions with pathogens.

## 1 Introduction

Communities of microbes associated with multicellular organisms, also known as microbiomes, can play a significant role in the health and disease of their hosts (Cho and Blaser, 2012; Lee and Hase, 2014; Oever and Netea, 2014; Robinson et al., 2010; Zilber-Rosenberg and Rosenberg, 2008). We are interested here in the skin-associated microbiome of frogs and the roles it may play in frog health. In general, the skin microbiome composition and structure in animals can influence host health by contributing to immune defenses and maintaining skin homeostasis (Sanford and Gallo, 2013). In amphibians, one key role of the skin microbiome is that it can serve as a primary defense mechanism against invading pathogens (Walke and Belden, 2016). The role of the amphibian skin microbiome in pathogen defense has become of great interest recently due to the global spread of the pathogen *Batrachochytrium dendrobatidis* (*Bd*), a chytrid fungus that causes the disease chytridiomycosis, which has led to dramatic declines and species extinctions in amphibians around the world (Fisher et al., 2009; Fisher and Garner, 2020; Scheele et al., 2019; Skerratt et al., 2007).

*Bd* infects keratinized epidermal cells, disrupting host osmoregulation and electrolyte balance, and often leading to mortality (Berger et al., 1999, 1998; Voyles et al., 2009). Interestingly, susceptibility to *Bd* varies widely among amphibian species, populations, and individuals (Jiménez and Sommer, 2017; Rosenblum et al., 2010). While this variation is influenced by multiple factors, including host genetics and environmental conditions, the skin-associated microbiome may also play a crucial role in determining susceptibility (Becker et al., 2015a). Studies have shown that several amphibian skin microbes can inhibit *Bd*, and that the structure of the skin microbiome can predict the severity of infection and disease outcomes (Bates et al., 2018; Becker et al., 2015a, 2015b; Harris et al., 2009a; Jani et al., 2017; Woodhams et al., 2007b, 2015). These findings have spurred interest in probiotics for amphibians, although results have been mixed (Becker et al., 2009; Harris et al., 2009a, 2009b; Kueneman et al., 2016a; Becker et al., 2021, 2015a, 2011; Knapp et al., 2022; Woodhams et al., 2020, 2012). Probiotic effectiveness often depends on the ability of beneficial bacteria to persist on the skin, which is influenced by the existing microbial community (Becker et al., 2015a; Knapp et al., 2022; Woodhams et al., 2012). This underscores the need for a deeper understanding of skin microbiome complexity and dynamics in amphibians (Becker et al., 2011; Costello et al., 2012; Garner et al., 2016).

*Bd* infections in frogs are primarily limited to the ventral skin surfaces and toes (Berger et al., 2005, 1998; North and Alford, 2008; Pessier et al., 1999). This pattern of infection may indicate a difference in the microbial communities and niche space available in certain regions of the body, warranting an examination of the microbiomes of different body regions to understand potential regional defenses against *Bd*. Evidence indicates that frog skin selects for specific microbes from the environment (Bates et al., 2018; Loudon et al., 2016; Walke et al., 2014), but whether there is selection for different microbes in body regions preferentially infected by *Bd* has not been examined.

Few studies have examined individual variability or within-individual variability in amphibian microbiomes, although such variations have been documented in humans and other animals (Asangba et al., 2022; Bouslimani et al., 2015; Grice et al., 2009; Krog et al., 2022; Shibagaki et al., 2017; Sugden et al., 2021). One study that examined individual variation in amphibian microbiomes over time found that skin microbiomes can vary between wild-captured individuals within the same population (Ellison et al., 2021). Furthermore, heterogeneity in microbiome structure among body regions has been detected in certain amphibian species (Bataille et al., 2016; Sabino-Pinto et al., 2016; Sanchez et al., 2017), suggesting that for at least some species, different skin regions may harbor distinct microbial communities.

In this study, we utilized high-throughput sequencing of bacterial 16S rRNA gene amplicons to characterize the skin microbiome of captive adult Sierra Nevada yellow-legged frogs (*Rana sierrae*) among and within individuals and their tank environments. This species has experienced dramatic population declines due to invasive fish and disease (Vredenburg et al., 2010, 2007). Restoration efforts for this species often involve head-starting, where frogs are reared to adulthood in captivity before being reintroduced into the wild. Captivity is known to alter the amphibian skin microbiome, with several studies finding differences in microbiome structure and diversity between captive and wild individuals across many amphibian species, likely due to environmental and dietary differences (Antwis et al., 2014; Becker et al., 2014; Kueneman et al., 2022; Loudon et al., 2014; Sabino-Pinto et al., 2016). These captivity-induced shifts in the microbiome could impact the success of reintroduction programs, warranting closer attention to the microbiome in captivity prior to release (Redford et al., 2012). Additionally, variability in the microbiome within populations or within individuals could affect health outcomes post-release and contribute to differences in *Bd* susceptibility and infection intensities observed within populations (Ellison et al., 2019; Jani and Briggs, 2014; Jiménez and Sommer, 2017; Rosenblum et al., 2010).

By examining the skin microbiome in a captive-reared population of *R. sierrae*, we sought to address the following questions: (1) How do captive *R. sierrae* skin microbiomes differ from their tank environment microbiome? (2) How much variation is there among microbiomes of frog individuals? (3) How much variation is there among microbiomes of different body regions within individuals? and (4) Are there consistent differences in the skin microbiome that correspond to body regions preferentially infected by *Bd*? We hypothesized that we would detect differences between frogs and their tank environments (Bataille et al., 2016; Walke et al., 2014), among individuals (Ellison et al., 2021), and among body regions (Bataille et al., 2016; Sabino-Pinto et al., 2016; Sanchez et al., 2017). Further, we hypothesized that certain microbes would be differentially abundant between body regions that tend to harbor *Bd* infections (ventral surfaces and feet) and body regions where infection is often absent (dorsal surfaces like the back).

## 2 Materials and Methods

### 2.1 Ethics statement

Non-invasive sampling of *Rana sierrae* individuals housed at the San Francisco Zoo was conducted with approval from the UC Davis IACUC (Protocol #18732) and the San Francisco Zoo Research Review Committee.

### 2.2 Frog population and handling

Frogs sampled for this study were reared to adulthood at the San Francisco Zoo from egg masses collected at a population in the Sierra Nevada Mountains located in the Desolation Wilderness (El Dorado County, California; ∼2500 m elevation). Adults, *i.e.*, those with snout–vent length (SVL) ≥ 40 mm, were tagged with 8 mm unique passive integrated transponder (PIT) tags, which allow for differentiation among individuals. Frogs were housed in tanks (groups of 8-13 individuals) filled with tap water purified using biological filters to remove toxic nitrogenous compounds and supplemented with Kent Marine R/O Right, a formulation of dissolved solids and electrolytes used to restore natural water chemistry to water that has been distilled, deionized, or purified by reverse osmosis.

### 2.3 Sample collection

We collected samples for this study from adult frogs and their tank environments. We wore nitrile gloves during sample collection from frogs and surfaces in tanks using sterile synthetic fine tip dry swabs (Medical Wire & Equipment, MW113). Prior to sampling, we rinsed each frog individual with 60 mL of sterile water (Culp et al., 2007; Lauer et al., 2007). From each frog, we collected a separate swab from each of the following body regions: back, outer hindlimbs, snout, vocal sack, ventral abdomen, inner forelimbs, forefeet, inner hindlimbs, hindfeet, and cloaca (Figure 1). Body regions were swabbed by taking 10 strokes to standardize sample material from regions of various sizes. For each frog, we also recorded the sex, tank identity, and individual identity (recording both the unique PIT tag number and “Tahoe ID” assigned to each frog by the Zoo). We collected swabs of surfaces in tanks including rock perches (above the water surface; two samples per tank), underwater rocks (rock perch submerged in water; one sample per tank), and tank walls (above the water surface; three samples per tank) by taking 40 strokes across each surface. Tank water was sampled by filling a 60 mL syringe, passing the water through a 0.22 μm Sterivex filter (Millipore), and repeating this process four times (total water filtered = 240 mL per sample; two filter samples collected per tank) (Ellison et al., 2019). All samples were kept on dry ice during collection and transferred to a -80 °C freezer for storage on the same day.

**Figure 1.**
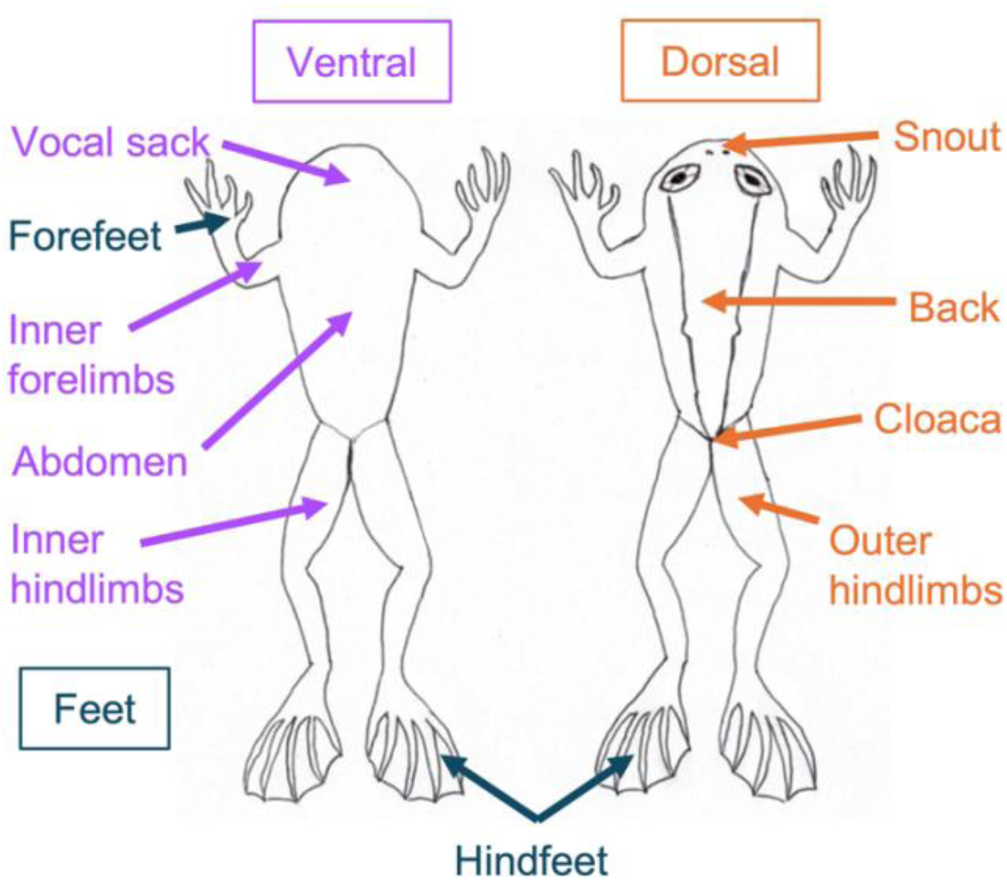
Diagram of *Rana sierrae* body regions sampled in this study. Ventral surfaces sampled are indicated with purple text and arrows; dorsal surfaces sampled are indicated with orange text and arrows; feet sampled are indicated with blue text and arrows. For limbs and feet, both the right and left were sampled. Forefeet were only sampled on the ventral surface, whereas hindfeet samples were collected from both the ventral and dorsal surfaces.

In this study, we analyzed samples collected from nine frogs (n = 90 microbiome swabs) and their tank environments (n = 18 microbiome swabs; n = 6 water filters) on July 28, 2015. These frogs were not distributed among tanks in a balanced manner: six frogs were co-housed in one tank, two frogs were co-housed in a second tank, and one frog was housed in a third tank.

### 2.4 DNA extraction

We extracted DNA from microbiome swabs and water filter samples using MoBio PowerSoil DNA Isolation Kits, using a modified protocol for low-biomass samples discussed with the manufacturer. Swabs or filters were swirled in the PowerBead tubes and left inside these tubes. Modifications to the manufacturer’s standard protocol included the following: (1) after adding Solution C1 and vortexing to mix, tubes were incubated at 65 °C for 10 minutes; (2) tubes were then secured in a bead beater set to “homogenize,” and bead-beated for a total of 3 minutes (90 seconds on, 60 seconds rest, followed by 90 seconds on); (3) all centrifugation steps throughout were done for 1 minute at 13,000 x *g* unless otherwise noted below; (4) we combined steps for Solutions C2 and C3 by adding 100 μL of each at once prior to 5 minute incubation on ice, (5) after the C2/C3 step, we transferred 700 μL of lysate to a clean collection tube and added 700 μL of Solution C4 and 600 μL of 100% ethanol before loading on the spin filters; (6) before washing the filter with solution C5, we inserted a step to wash with 650 μL of 100% ethanol; (7) After washing with solution C5, we dried the spin column by centrifuging for 2 minutes at 13,000 x *g*; (8) We added 60 μL of Solution C6 (heated to 60 °C) to the filter membrane, and we allowed this solution to sit on the filter for 5 minutes before centrifuging into a storage tube. Following DNA extraction, we quantified DNA concentration using a Qubit (Invitrogen, Carlsbad, CA, United States) and the dsDNA High Sensitivity Kit, and stored DNA extracts at -80 °C.

### 2.5 Sequence generation

Sequencing libraries were prepared following the protocol “16S Metagenomic Sequencing Library Preparation” (Part # 15044223 Rev. B, Illumina, Inc., San Diego, CA, USA) with some modifications. Briefly, we PCR amplified the hypervariable V3-V4 region of the bacterial 16S rRNA gene using 341F and 805R primers (Klindworth et al., 2013) with overhang adaptors (forward primer with overhang = 5’ TCGTCGGCAGCGTCAGATGTGTATAAGAGACAGCCTACGGGNGGCWGCAG; reverse primer with overhang = 5’ GTCTCGTGGGCTCGGAGATGTGTATAAGAGACAGGACTACHVGGGTATCTAATCC) from each sample in triplicate, using 4 uL DNA extract per reaction. We pooled PCR products from each sample (75 μL pool) and purified them using magnetic beads (Axygen AxyPrep Mag PCR Clean-Up Kit) using 60 μL beads per pool (for a 0.8X ratio of beads to PCR product), eluting in 25 μL of 10 mM Tris pH 8.5. Next, we attached dual indices and Illumina sequencing adaptors in a second round of PCR described in the Illumina protocol. We purified and normalized 25 μL of each index PCR product using SequalPrep Normalization Plate Kits (Invitrogen), following the manufacturer’s protocol with an extension of the binding step incubation to 2-6 hours. We then pooled 10 μL of purified, normalized, and indexed PCR product per sample, and used the Zymo Clean and Concentrator Kit to increase the DNA concentration of the pool following the manufacturer’s protocol (using a ratio of 5:1 of DNA Binding Buffer:PCR product, and final elution using 200 μL DNA Elution Buffer). We quantified DNA in the final pool using a Qubit (Invitrogen) and sent the pooled libraries to the UC Davis Genome Center DNA Technology Core for sequencing on an Illumina MiSeq (Illumina, Inc., San Diego, CA, USA) with v3 chemistry in 2 × 300 bp run mode.

We sequenced 20 negative control samples in addition to the true samples. These included six controls for swab sample collection (three dry swabs and three swabs rinsed with sterile water, processed in the same way as true samples), eight blank DNA extraction kit controls (two to three preparations from each of three PowerSoil kits for which no sample was added to the PowerBead tube, but otherwise processed in the same way as true samples), and six PCR controls (for which no sample DNA was added to the first PCR step, and subsequent processing was the same as for true samples).

### 2.6 Sequence processing

To demultiplex the sequence data, we used a modified version of a custom script designed by G. Jospin (https://github.com/gjospin/scripts/blob/master/Demul_trim_prep.pl). Primers were removed using cutadapt v. 3.5 (Martin, 2011) with Python v. 3.9.10 (van Rossum and Drake, 2009), discarding reads for which primers were not present. We processed the resulting sequences using the DADA2 v.1.24.0 (Callahan et al., 2016) workflow in R v. 4.2.1 with RStudio v. 2022.07.0-548 (Posit Team, 2022; R Core Team, 2022). We trimmed forward and reverse reads at 250 base pairs, truncated at the first quality score of 2, and removed them if the expected errors were greater than 4 (this removed 32.8% of sequences). We then merged reads and inferred amplicon sequence variants (ASVs; 7.9% of sequences did not pass through these steps). Next, we identified 1.8% of merged reads to be chimeric and removed them. After chimera removal, samples had a mean read depth of 23,832 with a range of 294 to 98,808 reads. We assigned taxonomy to genus level using the Ribosomal Database Project (RDP) naive Bayesian classifier algorithm and the SILVA high quality ribosomal RNA database v. 132, and species level assignments were made based on exact matching of ASVs to reference strains in the SILVA database (Quast et al., 2012; Wang et al., 2007; Yilmaz et al., 2014).

We then assigned unique names to ASVs, beginning with “SV” (sequence variant) followed by a number (e.g. SV1, SV2, etc.). We removed ASVs based on taxonomic classifications that were (1) non-bacterial at the domain level (including Eukaryota, Archaea, and those unclassified to domain), (2) chloroplasts, and (3) mitochondria, which resulted in 2,373 unique ASVs in the dataset.

We used Decontam v. 1.16.0 to identify putative contaminants, implementing the prevalence method with a probability threshold of 0.5 (which identifies sequences that have a higher prevalence in negative controls than in true samples) and setting the batch argument so that contaminants were identified independently within groups of samples associated with specific negative controls (Davis et al., 2018). We identified contaminants separately for each of the following four control sample groups: (1) dry swabs (n = 3) setting batch by the sample material so as to identify contaminants in swab samples and not filters, (2) swabs rinsed with sterile water (n = 3) setting batch by whether sampling involved rinsing with sterile water so as to identify contaminants from frogs that were rinsed prior to swabbing, (3) extraction kit blanks (n = 8) setting batch by the PowerSoil kit used so as to identify contaminants from each kit separately, and (4) PCR negative controls (n = 6) without specifying batch so as to identify contaminants associated with PCR across all samples. We then compiled a list of putative contaminants identified using each control group (102 unique ASVs) and removed them, leaving 2,271 unique ASVs in the dataset.

There has been an ongoing debate in the literature regarding the validity of using rarefaction as a sample normalization technique for microbiome data (Cameron et al., 2021; Gloor et al., 2017; McKnight et al., 2019; McMurdie and Holmes, 2014; Weiss et al., 2017). We chose to implement this method for many analyses because we were interested in community level comparisons that can become distorted using other normalization methods (McKnight et al., 2019). Additionally, rarefaction was shown to be more effective than other methods at controlling effects of sample library size when sample depths are very uneven (Schloss, 2023; Weiss et al., 2017), which is the case for our dataset. Therefore, for all subsequent sequence analyses except for DESeq2 differential abundance testing (see below), samples were rarefied at an even sampling depth of 2,727, which was the minimum depth of a true sample. All negative control samples had fewer than 2,727 reads and were therefore discarded at this step.

After rarefying the dataset, we aligned remaining sequences using DECIPHER v. 2.22.0 (Wright, 2016) and built a maximum likelihood tree with a GTR + Γ(4) + I model using phangorn v. 2.8.1 (Schliep et al., 2017; Schliep, 2011) on the UC Davis Bioinformatics Core High Performance Computing Cluster in R v. 4.1.0 (R Core Team, 2022). We midpoint rooted the tree using phangorn v. 2.9.0 (Schliep et al., 2017; Schliep, 2011).

The resulting dataset analyzed for this study included 1861 unique ASVs across 114 true samples.

### 2.7 Microbial sequence analysis and visualization

#### 2.7.1 Alpha diversity analysis

We considered two metrics of within-sample microbial community diversity (*i.e.* alpha diversity): observed richness and Shannon diversity. We calculated these metrics using the estimate_richness function in phyloseq v. 1.40.0 (McMurdie and Holmes, 2013). Shapiro-Wilk normality tests for groups of alpha diversity estimates that we sought to compare revealed that estimates for at least one group in each comparison were not normally distributed, warranting use of nonparametric statistical tests. We therefore implemented Kruskal-Wallis rank sum tests for significant differences in alpha diversity values between metadata groupings of the samples (including frog vs. environmental sample type, frog individual, and frog body region) using the kruskal.test function in base R v. 4.2.1 (R Core Team, 2022). For significant Kruskal-Wallis results (p ≤ 0.05), we performed *post hoc* Dunn tests with a Benjamini-Hochberg correction to control the false discovery rate (FDR) with multiple comparisons (dunnTest function in FSA v. 0.9.3 (Ogle et al., 2022)). Because individual identity was confounded with tank identity for several *post hoc* comparisons of frog individuals, we compared the percent of pairwise comparisons of co-housed individuals (*i.e.,* within tanks) that were significantly different to the percent of pairwise comparisons of individuals from distinct tanks (*i.e.,* between tanks) that were significantly different as a proxy of the relative importance of individual identity and tank identity to microbial alpha diversity.

#### 2.7.2 Beta diversity analysis

To assess community structure, we compared between-sample diversity (*i.e.* Beta diversity) using three ecological distance metrics: unweighted Unifrac, weighted Unifrac, and Bray-Curtis dissimilarities. We used the ordinate function in phyloseq to calculate these distances for different subsets of the data, and visualized ordinations using principal coordinate analysis (PCoA).

To test for significant differences in community structure between metadata groupings of the samples, we ran permutational multivariate analysis of variance (PERMANOVA) tests using the adonis function in vegan v. 2.6.2 with 9,999 permutations (Anderson, 2001; Oksanen et al., 2022). For each ecological distance metric, we ran a PERMANOVA on the whole dataset (frog samples and environmental samples) to test for significant differences in microbiome structure between frogs and environmental sample types, and then ran a PERMANOVA on the frog samples alone to test for significant differences among frog individuals and among frog body regions, as well as to test for significant effects of other metadata factors like frog sex (*i.e.,* males versus females). Next, for factors that rejected the null hypothesis in these PERMANOVA tests (p ≤ 0.05), we performed *post hoc* pairwise PERMANOVA tests to identify which levels within factors differed significantly, using the adonis.pair function in EcolUtils v. 0.1 with 9999 permutations (Salazar, 2022). We corrected p-values for multiple comparisons using the Benjamini-Hochberg procedure. We also calculated mean dispersion for factors included in PERMANOVAs and tested for significant differences using the betadisper and permutest.betadisper functions in vegan. For significant comparisons (p ≤ 0.05), we implemented *post hoc* Tukey honest significant differences tests to identify which levels within factors differed significantly in their group dispersion (using the TukeyHSD function in base R, which corrects p-values for the family wise error rate in multiple comparisons).

As we had done for alpha diversity, we assessed the relative importance of tank effects and individual effects to community structure based on *post hoc* pairwise comparisons of frog individuals, comparing the percent of significantly different comparisons for co-housed individuals to the percent of significantly different comparisons for individuals housed in distinct tanks

#### 2.7.3 Analysis of prevalence and relative abundance of taxa

We calculated the proportion of taxa shared between frog samples and environmental samples in the rarefied dataset. To do this, we first merged samples by frog and environment sample categories (*i.e.,* collapsing read counts within each sample category), and removed any ASVs that had zero counts across frog and environmental samples. Next, we calculated the proportion of ASVs that were present in both frog and environment samples (“shared”). We also calculated the proportion of bacterial families that were shared between frog and environmental samples by collapsing ASVs from the same families using the tax_glom function in phyloseq (specifying NArm=FALSE to include unclassified taxa) prior to merging samples by frog and environment sample categories.

We visualized and compared the relative abundance of taxa for the whole dataset (frog samples and environmental samples) and for a subset of the dataset (frog samples only). To do this, we first transformed the rarefied sample counts to relative abundances. To examine relative abundance of bacterial families across the whole dataset, we merged ASVs from the same families using the tax_glom function in phyloseq (specifying NArm=FALSE to include unclassified taxa). For visualization purposes, we filtered families to retain only those with mean relative abundance greater than 0.6%. We grouped the filtered families by a metadata factor to compare frog samples to the four environmental sample types and by taxonomic ranks desired for visualization and then plotted mean relative abundance with standard error bars faceted by levels of the metadata grouping factor. For visualizations of the relative abundance of taxa within frog samples, we focused on two dominant ASVs. We used the prune_taxa function in phyloseq to retain only the ASVs of interest, grouped data by metadata factors (frog individual or frog body region) and taxonomic rank for visualization, and plotted the mean relative abundances of these ASVs with standard error bars for each metadata factor (individual and body region).

To determine whether the relative abundance of bacterial families or ASVs differed significantly between metadata groupings, we implemented nonparametric Kruskal-Wallis rank sum tests followed by *post hoc* Dunn tests when results of Kruskal-Wallis tests were significant (p ≤ 0.05). We corrected p-values from Dunn tests using the Benjamini-Hochberg procedure. For *post hoc* comparisons of frog individuals, we compared the percent of pairwise comparisons within tanks that were significant to the percent of pairwise comparisons between tanks that were significant, again to assess the relative importance of individual identity and tank identity.

For frog samples, we determined whether the relative abundances of two dominant ASVs of interest were significantly different from each other within sample metadata groupings (*i.e.,* within each individual or within each body region). To do this, we used nonparametric Wilcoxon signed rank exact tests with the Benjamini-Hochberg procedure for p-value correction.

#### 2.7.4 DESeq2 differential relative abundance testing

We used DESeq2 v. 1.36.0 in R on filtered but un-rarified merged read counts to determine which ASVs showed significant log2 fold differences between frog body regions (Love et al., 2014). Based on previous findings that *Bd* infection is more prominent on ventral surfaces and toes than on the dorsal back surface of many anurans (Berger et al., 2005, 1998; North and Alford, 2008; Pessier et al., 1999), we chose to compare frog back samples to (1) abdomen samples, (2) inner hindlimb samples, (3) hind feet samples, and (4) forefeet samples, to examine whether bacterial community members were differentially associated with these body regions.

First, we used the phyloseq_to_deseq2 function to format the raw read count data from frog samples for DESeq2. Our ASV counts table was sparse, with only two ASVs present across all samples. Therefore, to prevent geometric means and estimated size factors for DESeq2 sample normalization from being influenced solely by these ASVs, we calculated geometric means across samples for each ASV by ignoring samples with zero counts. Then, we estimated size factors for each sample based on the geometric means (applying the estimateSizeFactors function). We filtered out low relative abundance ASVs with 10 or fewer reads total across frog samples (this filtered out 1984 unique ASVs, leaving 287 unique ASVs across the frog samples). We then ran the DESeq function on the dataset for each contrast of interest, identifying ASVs that showed significant log2 fold differences (*i.e.,* that showed differential relative abundance; Benjamini-Hochberg corrected p-values ≤ 0.05). For ASVs that showed differential relative abundance based on DESeq2 normalized counts, we calculated the mean relative abundance across body regions from the rarefied dataset and implemented Kruskal-Wallis rank sum tests to determine whether mean relative abundance also differed significantly among body regions.

#### 2.7.5 *Bd*-inhibitory predictions for select taxa

We predicted putative *Bd*-inhibitory function of microbial community members of interest. Predictions were based on a database of full length 16S rRNA gene sequences from bacteria that were isolated and assayed for their effects on growth of *Batrachochytrium* pathogens (Woodhams et al., 2015). This database, which is regularly updated, has been used in several previous studies to predict anti-*Bd* function from amplicon data (Bletz et al., 2017; Chen et al., 2022; Jiménez et al., 2022; Kueneman et al., 2016a, 2022; Muletz Wolz et al., 2018). We used the strict inhibitory subset of the database (AmphiBac_InhibitoryStrict_2023.2; accessed from https://github.com/AmphiBac/AmphiBac-Database/) which included sequences from 2,056 inhibitory taxa. We used NCBI nucleotide BLAST to make a multiple sequence alignment with ASV sequences as queries and the inhibitory database as subject sequences (Camacho et al., 2009). We checked that query coverage was 100%, and then documented cases where the percent identity was ≥99% (*i.e.,* cases where the query ASVs shared ≥99% sequence similarity with an inhibitory database sequence).

## 3 Results

### 3.1 Alpha diversity

We used two alpha diversity metrics to evaluate within-sample diversity: the observed richness (*i.e.,* the number of ASVs in the rarefied dataset) and Shannon diversity, which incorporates ASV richness and relative abundance (*i.e.,* richness and evenness).

#### 3.1.1 Frogs vs. tank environment microbiome diversity

Within-sample diversity was significantly lower in frog samples than in tank environment sample types (including rock perch, tank wall, tank water, and underwater rock samples) based on both the observed richness (Kruskal-Wallis, chi-squared = 55.62, df = 4, p < 0.001; Dunn test, p < 0.05; Table S1; Figure 2A) and Shannon diversity (Kruskal-Wallis, chi-squared = 56.77, df = 4, p < 0.001; Dunn test, p < 0.05; Table S1; Figure 2B).

**Figure 2.**
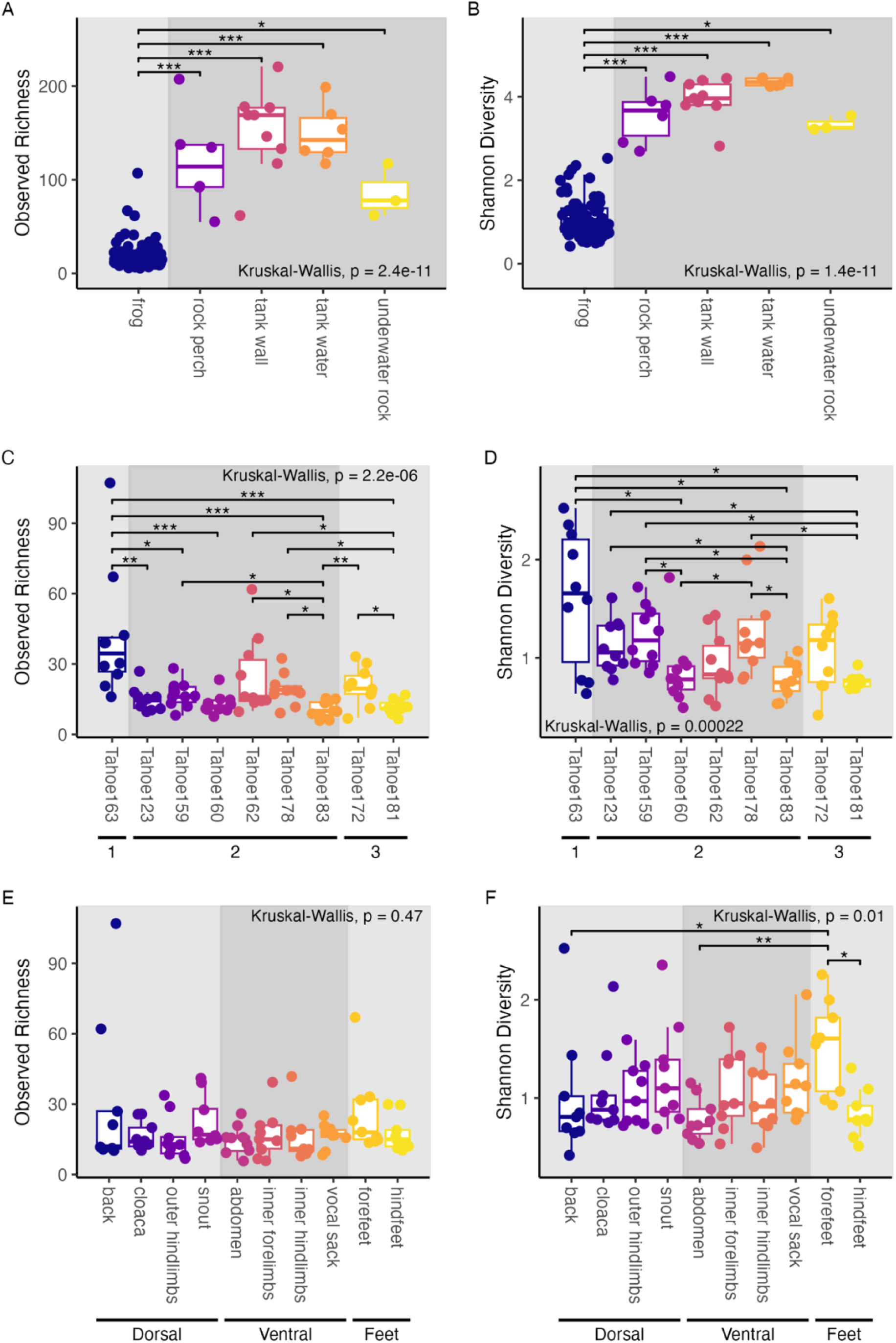
Alpha diversity based comparisons of sample groupings. Within-sample diversity in terms of observed richness **(A,C,D)** and Shannon diversity **(B,D,F)**; **(A-B)** Comparison of frog and tank samples; Boxplots and points are colored by the type of sample substrate and panel background shading differentiates frog samples from tank environment samples; **(C-D)** Comparison of samples from distinct frog individuals; Boxplots and points are colored by frog individual ID; panel background shading differentiates groups of individuals housed in separate tank aquaria (1, 2, and 3); **(E-F)** Comparison of samples from distinct frog body regions; Boxplots and points are colored by frog body region; panel background shading differentiates groups of body regions sampled (dorsal, ventral, and feet); **(A-F)** Results of Kruskal-Wallis tests are shown; *post hoc* Dunn test results are displayed as significance bars where applicable (“*” = p ≤ 0.05; “**” = p ≤ 0.01; “***” = p ≤ 0.001).

#### 3.1.2 Variation in frog microbiome diversity by individual and tank identity

Within frog samples, we found that both observed richness and Shannon diversity differed significantly by frog individual (Observed: Kruskal-Wallis, chi-squared = 40.84, df = 8, p < 0.001; Shannon: Kruskal-Wallis, chi-squared = 29.95, df = 8, p < 0.001; Figure 2C, D). For observed richness, we identified 12 significant differences out of 36 pairwise comparisons of our 9 individuals (Dunn test, p < 0.05; 33% of comparisons were significant; Table S2). While 25% of within-tank pairwise comparisons of individuals were significantly different, 40% of between-tank pairwise comparisons of individuals were significantly different. For Shannon diversity, we identified 11 significant differences out of 36 pairwise comparisons of individuals (Dunn test, p < 0.05; 30.5% of comparisons were significant; Table S2). 31.3% of within-tank pairwise comparisons were significantly different and 30% of between-tank pairwise comparisons were significantly different. These results show that there was variability in the number of ASVs and their evenness among frog individuals and among tanks, but that a greater percentage of differences in observed richness were between tanks than among individuals within tanks.

#### 3.1.3 Variation in frog microbiome diversity by body region

We found that Shannon diversity differed significantly by frog body region, (Kruskal-Wallis, chi-squared = 21.54, df = 9, p = 0.010; Figure 2F), but that observed richness did not (Kruskal-Wallis, chi-squared = 8.67, df = 9, p = 0.47; Figure 2E). Frog forefeet harbored higher Shannon diversity than the abdomen, back, and hind-feet in *post hoc* comparisons (Dunn tests, p < 0.05; Table S3). In other words, while all frog body regions harbored a similar number of microbial community members, the forefeet harbored communities with higher evenness than certain other regions.

### 3.2 Beta diversity

We evaluated differences in community structure (*i.e.,* Beta diversity) using three metrics. Unweighted UniFrac distance is calculated from the community phylogenetic tree as the unique fraction of branch length within a sample community that is not shared with other communities sampled (Lozupone and Knight, 2005). This metric, therefore, can be thought of as measuring distances based on community membership (presence/absence). Weighted UniFrac takes into account the relative abundances (*i.e.,* evenness) of branch lengths in addition to membership, giving more weight to dominant organisms than rare ones (Lozupone et al., 2007). Bray-Curtis also takes into account species richness and relative abundances, but this metric is not informed by phylogeny.

#### 3.2.1 Frog vs. tank environment microbiome structure

Microbial community structure was significantly different between frog samples and tank environment sample types based on all three ecological distance metrics (PERMANOVA, p < 0.001; Figure 3A-C, Table S4). *Post hoc* pairwise PERMANOVA tests revealed that all groups (frog, tank water, tank wall, rock perch, and underwater rock) were significantly different from each other based on all three metrics (PERMANOVA, p < 0.05; Table S5). However, the relative importance of community characteristics measured by each metric differed. We found that differences between microbial assemblages on frogs and those from environmental samples explained the highest amount of variation in weighted Unifrac (69%), followed by Bray-Curtis (45%), and finally unweighted Unifrac (29%) (Table S4). In other words, differences in the relative abundances of community members explained a higher proportion of variation between frogs and their environment than presence/absence of community members alone.

**Figure 3.**
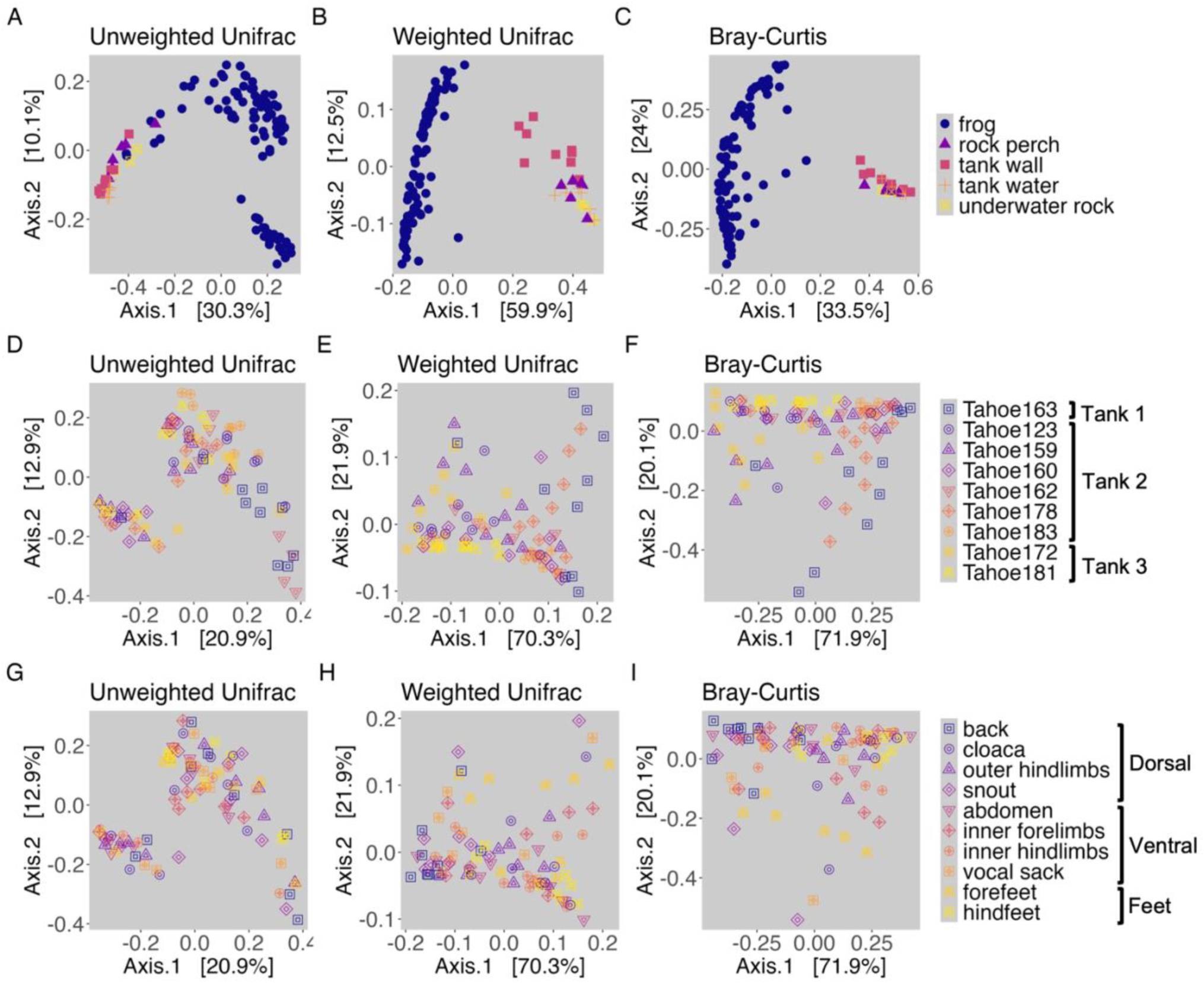
Beta diversity based comparisons of sample groupings. **(A-C)** Microbial community structure of frogs and environment sample types (rock perch, tank wall, tank water, and underwater rock); points are colored and shaped by frog category and environmental sample types; **(D-F)** Microbial community structure of frog individuals; points are colored and shaped by frog individual (i.e., unique ID); **(G-I)** Microbial community structure of frog body regions; points are colored and shaped by frog body region; **(A, D, G)** PCoA visualizations of unweighted Unifrac distances; **(B, E, H)** PCoA visualizations of weighted Unifrac distances; **(C, F, I)** PCoA visualizations of Bray-Curtis dissimilarities.

PERMANOVA tests are sensitive to differences in group dispersion (*i.e.,* within-group variance) and thus significant effects detected by these tests could indicate differences in the average location of groups in ordination space (*i.e.,* group centroids), differences in group dispersion, or some combination of the two (Anderson, 2001). If PERMANOVA tests indicate a significant effect of a factor and group dispersions for that factor do not differ significantly, then we know that centroid location differs for these groups. However, if the group dispersions are significantly different, then our tests are unable to distinguish whether differences are due to dispersion alone or some combination of dispersion and centroid location. We found group dispersion between frogs and environment sample types did not differ significantly for unweighted Unifrac (betadisper permutest, p = 0.16; Table S6), indicating that differences based on phylogenetically informed community membership were due to differences in mean centroids. There were significant differences in group dispersion, however, for weighted Unifrac and Bray-Curtis (betadisper permutest, p < 0.001; Table S6). For weighted Unifrac, pairwise comparisons of group dispersion (for frog, rock perch, tank wall, tank water, and underwater rock sample groupings) revealed that five out of ten comparisons were significantly different, which included two out of the four comparisons with frog samples (TukeyHSD, p < 0.05; Table S7). For Bray-Curtis dissimilarity, three out of ten pairwise comparisons of group dispersion were significantly different, and these represented three of the four comparisons with frog samples (TukeyHSD, p < 0.05; Table S7). However, ordination visualizations (Figure 3A-C) showed that frog samples clustered separately from environmental sample types for all three measures of community structure, which is evidence that in cases where differences in dispersion were detected between frog and environment samples, both the group dispersions and centroid locations may have been distinct. We also note that these ordinations comparing frog and environmental samples (Figure 3A-C) show a characteristic “horseshoe effect,” which could be a consequence of saturation of the distance metrics (Morton et al., 2017). Since distance metrics cannot discriminate between samples that do not share ASVs, this saturation can occur when the dataset has sparse counts of ASVs across samples creating a “band table” (Morton et al., 2017). This effect does not alter our interpretation of the ordinations, and it actually provides further evidence of the high dissimilarity between environment and frog samples.

#### 3.2.2 Drivers of frog microbiome structure

Our PERMANOVA model to explain variation in community structure within frog samples revealed that the factor with largest effect was individual identity (PERMANOVA, p < 0.001; Table S8). This factor explained the highest amount of variation in weighted Unifrac distances (41%), followed by Bray-Curtis (37%) and lastly by unweighted Unifrac (22%). Even after accounting for the effects of individual identity, we also found a significant effect of the body region sampled (PERMANOVA, p < 0.05; Table S8). Body region explained the highest amount of variation in Bray-Curtis (32%), followed by weighted Unifrac (28%), and explained the least amount of variation in unweighted Unifrac distances (11%). Shuffling the order of factors (individual and body region) in the PERMANOVA model did not alter these results, and the interaction term for these factors was not significant and was dropped from the model. We also tested whether frog sex (*i.e.,* males versus females) explained variation in the microbiome. However, when this factor was included after individual identity in sequential PERMANOVA models, it was not significant; thus, sex was also dropped from the model.

Overall, the results of the omnibus model show that frog individual (which includes tank variation) and body region explained more variation in community structure when relative abundances of microbes were incorporated than when community membership was considered alone, which may indicate that shifts in abundant community members were more important to explaining differences than shifts in rare community members. Further, while differences among frog individuals were better explained by phylogenetic distance-based metrics, differences among body regions were better explained by a taxonomy-based metric.

#### 3.2.3 Variation in frog microbiome structure among individuals and tanks

To determine how many frog individuals were driving variation in community structure explained by this factor, we conducted *post hoc* pairwise PERMANOVA tests. We also compared the relative importance of individual identity to that of tank identity in pairwise comparisons, as individual identity was confounded with tank identity for many comparisons.

Although frog individual identity explained the smallest amount of variation in unweighted Unifrac distances overall compared to other metrics, that variation was driven by the highest number of significantly different individuals. Out of 36 possible combinations of frog individuals, 26 pairwise comparisons were significantly different for unweighted Unifrac (pairwise PERMANOVAs, p < 0.05; 72.2% of comparisons were significant; Table S9; Figure 3D). This included 68.8% of pairwise comparisons of co-housed individuals (*i.e.,* within-tank), and 75% of pairwise comparisons of individuals between distinct tanks. For weighted Unifrac, 19 of the 36 pairwise comparisons of individuals were significantly different (pairwise PERMANOVAs, p < 0.05; 52.8% of comparisons were significant; Table S9; Figure 3E). This included 25% of pairwise comparisons of co-housed individuals and 75% of pairwise comparisons of individuals between tanks. Finally, for Bray-Curtis dissimilarity, 18 of the 36 pairwise comparisons of individuals were significantly different (PERMANOVA, p < 0.05; 50% of comparisons were significant; Table S9; Figure 3F). This included 31.3% of pairwise comparisons of co-housed individuals within tanks and 65% of pairwise comparisons of individuals between tanks. In summary, differences in relative abundance-weighted community structure showed a greater association with tank identity than individual identity. Differences in community membership showed only a slightly stronger association to tank identity than individual identity, and individuals within tanks showed over two times as many pairwise differences in community membership than they did for relative abundance-weighted community structure.

We found no significant differences in group dispersion by individual for weighted Unifrac (betadisper permutest, p > 0.05; Table S10) or Bray-Curtis (betadisper permutest, p > 0.05; Table S10). In addition, we did not detect significant differences in dispersion among individuals in *post hoc* pairwise comparisons for unweighted Unifrac (TukeyHSD, p > 0.05; Table S11). These results confirm that significant differences in community structure described above represented differences in centroid locations of community structure rather than differences in group variances among frog individuals.

#### 3.2.4 Variation in frog microbiome structure among body regions

We used pairwise PERMANOVA tests to determine which frog body regions drove the variation in community structure explained by this factor. For unweighted Unifrac, none of the 45 pairwise comparisons of body regions were significantly different after correcting p-values for multiple comparisons (PERMANOVA, p > 0.05; Table S12; Figure 3G). For weighted Unifrac, nine pairwise comparisons of body regions were significantly different (PERMANOVA, p < 0.05; 20% of comparisons were significant; Table S12; Figure 3H), and these included seven significant differences between the back and other body regions, and two significant differences between the snout and other body regions (hindfeet and cloaca were significantly different from both the back and the snout; the abdomen, forefeet, inner forelimbs, inner hindlimbs, and outer hindlimbs were all significantly different from the back). For Bray Curtis, 13 pairwise comparisons of body regions were significantly different (PERMANOVA, p < 0.05; 28.9% of comparisons were significant; Table S12; Figure 3I), including the same seven significant differences with the back and two significant differences with the snout that were identified for weighted Unifrac, as well as four additional significant differences with the snout only identified for Bray-Curtis (the snout also differed from the abdomen, forefeet, inner hindlimbs and outer hindlimbs for Bray-Curtis). The vocal sack was the only region that never differed significantly from other body regions in terms of community structure.

Group dispersion by body region did not differ significantly for any distance metrics (betadisper permutest, p > 0.05; Table S13). These results indicate that significant results from pairwise PERMANOVA tests comparing frog body regions represented significant differences in group centroids for community structure and not differences in dispersion among body regions. Further, our results suggest that relative abundances of dominant community members rather than community membership explain differences among body regions.

### 3.3 Prevalence and relative abundance of taxa

#### 3.3.1 Frog vs. tank environment microbial taxa

To investigate the similarity in presence of taxa between frog samples and environmental samples, we calculated the proportion of taxa shared between frog and environment. We found that 15.4% of ASVs were shared between frog and environmental samples (212 out of 1,378 ASVs). We note that the proportion of shared ASVs will most likely be reduced compared to the proportion of shared OTUs, the unit of comparison in many previous studies (Kueneman et al 2013; Bates et al 2018; Walke et al 2014), because OTU calling involves collapsing sequence variants by a threshold of similarity (usually 97%), while ASVs are not clustered. We also looked at the proportion of families shared between frog and environmental samples, which was 38.4% (91 out of 237 bacterial families).

To better understand the composition of microbiomes defining different types of samples, we visualized taxonomic families that had mean relative abundance greater than 0.06% across all samples (an arbitrary threshold selected for optimized visualization). This further revealed that the distribution of taxa were distinct between frog samples and the four types of environmental samples (Figure 4). Frog-associated communities were dominated by the families Burkholderiaceae (phylum Proteobacteria; mean relative abundance of 48.0 ± 2.6% across frogs), Rubritaleaceae (phylum Verrucomicrobia; mean relative abundance of 39.5 ± 2.5% across frogs), and to a lesser extent the families Pseudomonadaceae and Alcanivoracaceae (both in phylum Proteobacteria; mean relative abundance of 7.3 ± 1.2% and 2.7 ± 0.2%, respectively). The environment-associated communities were, for the most part, made up of lower relative abundances (mean relative abundance <17%) of an increased number of families representing several phyla in addition to the Proteobacteria and Verrucomicrobia, including the Actinobacteria, Bacteroidetes, Deinococcus-Thermus, and Firmicutes (Figure 4).

**Figure 4.**
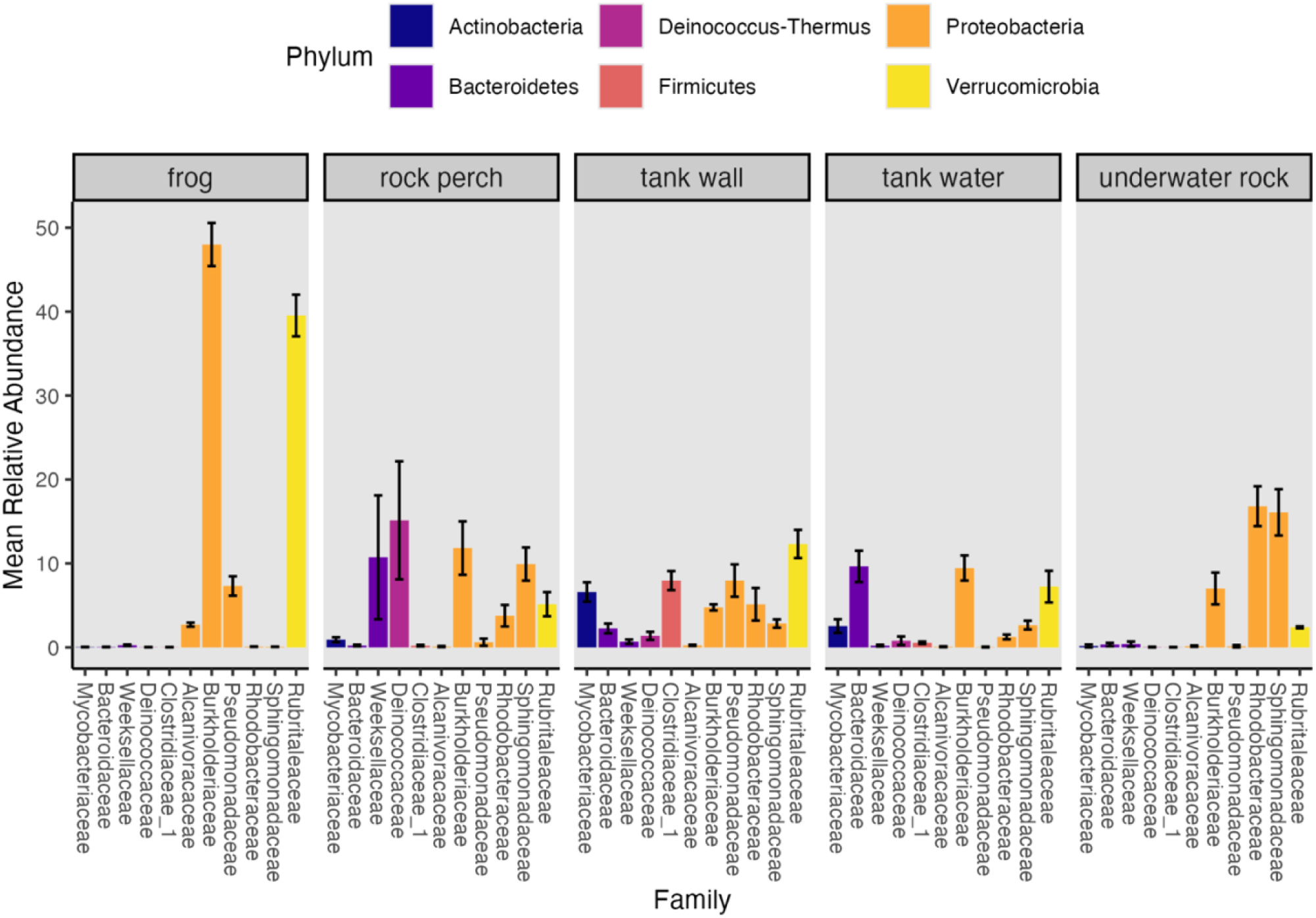
Mean relative abundances of top represented bacterial families. Bacterial families with mean relative abundance greater than 0.6% across samples are shown; families are colored and ordered by phylum; plot is faceted by frog, rock perch, tank wall, tank water, and underwater rock samples; error bars represent the standard error around the mean relative abundance.

Next, we examined whether there were significant differences in the relative abundance of taxa between frogs and environmental sample types. Relative abundance of the family Burkholderiaceae was significantly higher on frogs than on the four environment sample types (Kruskal-Wallis, chi-squared = 357.31, df = 4, p < 0.001; Dunn test, p < 0.001; Table S14; Figure 4). Within the Burkholderiaceae, one ASV (“SV2,” which was unclassified at the genus level) was dominant on frogs, making up 98.8% of Burkholderiaceae rarefied read counts across frog samples and 31.42% of Burkholderiaceae rarefied read counts across environmental samples. While SV2 was present in 100% of samples, its relative abundance was significantly different between frogs and environmental sample types (Kruskal-Wallis, chi-squared = 56.492, df = 4, p < 0.001), with significantly higher relative abundance on frogs than in the environmental sample types (Dunn tests, p < 0.01; Table S15).

The family Rubritaleaceae was also dominated by one ASV present in 100% of samples (“SV1,” which was unclassified at the genus level) that made up 99.9% of Rubritaleaceae rarefied read counts across frog samples and 92.68% of Rubritaleaceae rarefied read counts across environmental samples. While the relative abundance of the family Rubritaleaceae was not significantly different between frogs and environment sample types (Kruskal-Wallis, chi-squared = 3.6618, df = 4, p = 0.4537; Figure 4), SV1 relative abundance was significantly different between these groups (Kruskal-Wallis, chi-squared = 45.626, df = 4, p < 0.001), with higher relative abundance on frogs than in environmental samples (Dunn tests, p < 0.01; Table S15).

#### 3.3.2 Variation in dominant frog-associated taxa among individuals and tanks

We examined variation in the relative abundances of the two dominant ASVs on frogs, SV1 (family Rubritaleaceae) and SV2 (family Burkholderiaceae), across frog individuals (Figure 5A) to ascertain whether variation in these ASVs contributed to differences between individuals. The relative abundance of both SV1 and SV2 differed significantly by frog individual (Kruskal Wallis, SV1: chi-squared = 40.828, df = 8, p < 0.001, SV2: chi-squared = 27.863, df = 8, p < 0.001). *Post hoc* Dunn tests revealed that out of 36 pairwise comparisons of the nine frog individuals, SV1 relative abundance differed significantly in 13 comparisons (*i.e,* 36.1% of comparisons were significant; Dunn tests, p < 0.05; Table S16). Because frogs were not evenly distributed among distinct tank environments, we quantified the proportion of between-tank and within-tank individual comparisons that differed and found that 25% of between-tank individual comparisons and only 6.25% of within-tank individual comparisons were significantly different in terms of SV1 relative abundance. SV2 relative abundance differed significantly in six of the 36 pairwise comparisons of individuals (*i.e.,* 16.7% of comparisons were significant; Dunn tests, p < 0.05; Table S16). While 55% of between-tank pairwise comparisons of individuals were significant, only 12.5% of within-tank pairwise comparisons of individuals were significant. Thus, the relative abundances of these ASVs both appeared to be primarily associated with tank identity rather than individual identity. Further, ordering individuals by the mean relative abundance of SV1 also grouped them by their associated tank identity, providing support for the importance of tank identity to this taxon’s relative abundance (Figure 5A).

**Figure 5.**
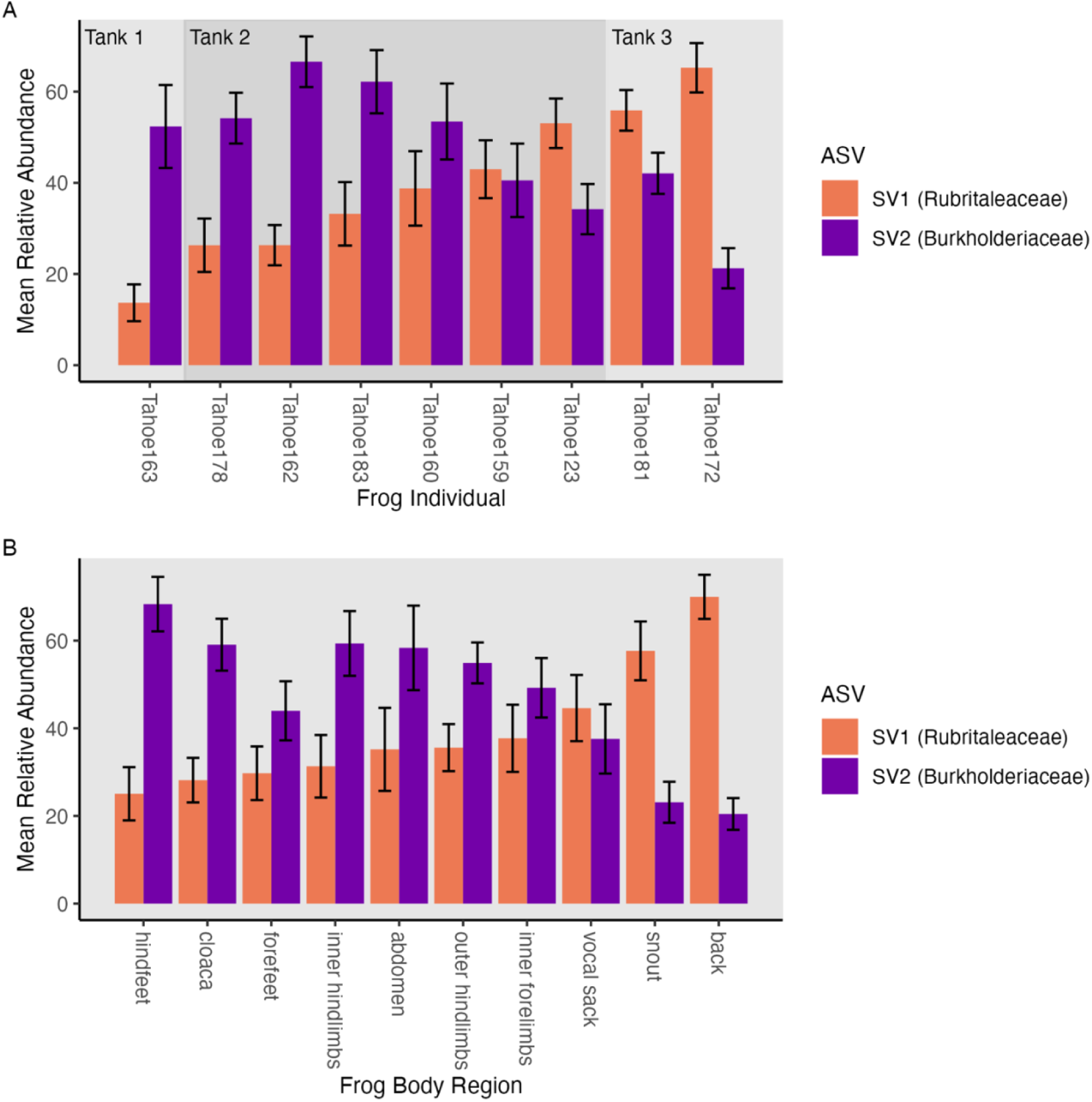
Mean relative abundances of two dominant frog-associated taxa. Mean relative abundances of two dominant ASVs (SV1, family Rubritaleaceae, and SV2, family Burkholderiaceae) on frogs; **(A)** Mean relative abundances by individual identity; panel background shading differentiates groups of individuals housed in distinct tank aquaria; **(B)** Mean relative abundances by body regions; **(A-B)** Relative abundances were calculated from the rarefied dataset; frog individuals and frog body regions are ordered from lowest to highest mean relative abundance of SV1; bars are colored by ASV and associated bacterial family; error bars represent the standard error around the mean relative abundance.

We also tested whether the relative abundance of SV1 was significantly different from the relative abundance of SV2 within each individual (*i.e.,* whether the relative abundance of one of these ASVs was consistently higher than the other). Wilcoxon signed rank test results revealed that SV1 and SV2 relative abundances were significantly different within three frog individuals, each of which was housed in a different tank. One individual had significantly higher SV1 relative abundance and two individuals had significantly higher SV2 relative abundance (Wilcoxon, p < 0.05; Table S17).

#### 3.3.3 Variation in dominant frog-associated taxa among body regions

The relative abundance of both SV1 and SV2 also differed significantly by frog body region (Kruskal-Wallis, SV1: chi-squared = 27.941, df = 9, p < 0.001, SV2: chi-squared = 35.697, df = 9, p < 0.001). Out of 45 pairwise comparisons of body regions, SV1 relative abundance differed significantly in nine comparisons. SV1 relative abundance was significantly higher on the back than on seven other body regions (the abdomen, cloaca, forefeet, hindfeet, inner forelimbs, inner hindlimbs, and outer hindlimbs; Dunn tests, p < 0.05; Table S18), and was significantly higher on the snout than on two other body regions (the cloaca and hindfeet; Dunn tests, p < 0.05; Table S18). SV2 relative abundance differed significantly in 12 out of 45 pairwise comparisons, with significantly lower relative abundance on the back than on six other body regions (the abdomen, cloaca, hind-feet, inner forelimbs, inner hindlimbs, and outer hindlimbs; Dunn tests, p < 0.05; Table S18), significantly lower relative abundance on the snout than on five other body regions (the abdomen, cloaca, hind-feet, inner hindlimb, and outer hindlimbs; Dunn tests, p < 0.05; Table S18), and significantly higher SV2 relative abundance on hindfeet than on the vocal sack (Dunn test, p = 0.029; Table S18).

We next examined whether the relative abundance of SV1 was significantly different from the relative abundance of SV2 within each body region. We found that the relative abundance of SV1 was significantly higher than that of SV2 on both the back and the snout (Wilcoxon, p < 0.05; Table S19).

### 3.4 DESeq2 Differential Abundance Testing

To identify ASVs that were differentially abundant between body regions known to experience higher *Bd* infection and pathogenesis (*e.g.,* ventral surfaces and toes) and body regions known to have markedly lower *Bd* infection (*e.g.,* dorsal surfaces, mainly the back) we implemented DESeq2 analysis on raw read count data. We individually compared DESeq2 normalized counts from the abdomen, the inner hindlimbs, the hindfeet, and the forefeet to the back. The analysis identified one unique ASV, an undescribed member of the family Burkholderiaceae, that showed significant log2 fold higher normalized counts on the abdomen compared to the back (SV56; estimated log2 fold difference of 24.18; Table S20). The mean relative abundance of this ASV in the rarefied dataset, however, was not significantly different among body regions (Kruskal Wallis, chi-squared = 5.5597, df = 9, p = 0.783; Figure S1).

### 3.5 *Bd*-Inhibitory Predictions

We found that the dominant undescribed Burkholderiaceae across frog samples (SV2), shared 100% sequence identity with a known *Bd*-inhibitory taxon, AmphiBac_1576 (Woodhams et al., 2015). Another undescribed Burkholderiaceae that showed significant log2 fold higher normalized relative abundance on the *R. sierrae* abdomen than on their back (SV56; *see above*), shared 99.53% sequence identity with the same inhibitory taxon. The dominant undescribed Rubritaleaceae across frog samples (SV1), did not share ≥99% sequence similarity with any taxa from the inhibitory database.

## 4 Discussion

Our fine-scale analysis of the skin microbiome identified characteristics that vary within and among frog individuals and their tank environments. While captive frogs harbored distinct microbial communities compared to their local tank environment, more variation in frog microbiomes was associated with distinct tank enclosure than with individual frog identity. In addition, there were detectable differences between microbiomes of body regions preferentially infected with *Bd* compared to those regions that infrequently experience infection. Further, elevated relative abundances of putatively *Bd*-inhibitory microbes were localized in body regions where we would expect interactions with *Bd* to occur. Together, these results help elucidate the captive microbiome of the endangered Sierra Nevada yellow-legged frog, *R. sierrae*, and provide a basis for predicting microbiome-pathogen interactions.

### 4.1 Frog skin microbiomes were distinct from their tank environment microbiome and were dominated by fewer organisms

We hypothesized that frog skin microbiomes would be distinct from their surrounding tank environment microbiome, which was supported by our results for within-sample community diversity (*i.e.,* alpha diversity), community structure (*i.e.,* Beta diversity), and presence and relative abundance of microbial community members. We found that tank substrates and water harbored significantly higher within-sample diversity than frogs (Figure 2A,B), which agrees with a previous study of wild *R. sierrae* showing that lake water communities had higher observed richness than frog associated communities (Ellison et al., 2019). However, this finding differed from previous evidence that lake water microbiomes had reduced or equal diversity compared to microbial communities associated with several other species of post-metamorphic amphibians (Bates et al., 2018; Kueneman et al., 2014). The discrepancy between our results here and those of these prior studies has multiple possible explanations. One possibility is that lake water collected by Ellison et al. (2019) and tank water collected here were unusually diverse compared to other environments. Another possibility (and we note, both could be occurring) is that *R. sierrae* may harbor lower diversity microbiomes than other species. Reduced diversity is linked to clinical signs of chytridiomycosis (Becker and Harris, 2010) while higher community richness has been shown to correlate with host persistence after *Bd* invasion (Jani et al., 2017). Therefore, if *R. sierrae* microbiomes harbor lower diversity and richness than other species, this could relate to their high susceptibility to *Bd* (Vredenburg et al., 2010). Additional studies that directly compare the diversity of *R. sierrae* microbiomes to other species would be useful here.

We also found that community structure significantly differed between frog-associated and environment-associated microbiomes (Figure 3A-C), which has been previously reported in studies of both captive and wild amphibian populations (Albecker et al., 2019; Bates et al., 2018; Fitzpatrick and Allison, 2014; Jani et al., 2017; Jani and Briggs, 2014; Kueneman et al., 2014; Walke et al., 2014). There are several factors that make amphibian skin a unique and complex environment that could lead to such differences. Mucosal secretions, anti-microbial peptides (AMPs), and other secretions produced by the host regulate microbial presence and abundance on the skin, as do other microbes and the anti-microbial metabolites they produce, all of which vary between and within host species (Lillywhite and Licht, 1975; Myers et al., 2012; Tennessen et al., 2009; Walke et al., 2014; Woodhams et al., 2010, 2007a, 2006a, 2006b). By affecting which microbes can exist and persist on the skin, these interacting skin components act as a filter for microbes from the environment.

Previous studies found variable proportions of taxa shared between amphibian- and environment-associated communities, and usually dominant microorganisms on amphibians were different from those in environmental assemblages (Bates et al., 2018; Kueneman et al., 2014; Walke et al., 2014). Further, abundant microorganisms on amphibians have been shown to be rare in the environment (Bates et al., 2018; Kueneman et al., 2014; Walke et al., 2014). Here, we found that only 15.4% of ASVs were shared between frogs and environmental samples. We also looked at the proportion of shared bacterial families between frog and environmental samples, which was higher at 38.4%. We note that the proportion of taxa shared with their environment may be lower for captive frogs than their wild counterparts, as was shown previously (Bataille et al., 2016).

Additionally, in our study, a major difference in the distribution of taxa between frogs and their tank environments was that the two sequence variants found to be dominant on frogs (SV1 in the family Rubritaleaceae and SV2 in the family Burkholderiaceae) both showed significantly lower relative abundance on tank and perch substrates and in tank water (Figure 4). This supports the idea that high relative abundances of these bacteria were selected for by the frog’s skin (Loudon et al., 2016; Walke et al., 2014). A caveat of these statistical comparisons is that the data used is compositional (*i.e.,* relative abundances must sum to 100%). Therefore, care must be taken with the interpretation of differences in relative abundances across samples since they do not represent absolute abundances and are standardized to the rarefied read count. For example, the higher relative abundances of these two taxa on frogs than in their environment could have resulted from higher absolute abundances on frogs, but it also could have resulted from reduced abundances of other taxa on frogs that inflated the relative abundance of these two ASVs. Regardless, it interesting that microbial relative abundances on *R. sierrae* skin were dominated by only two sequence variants, and this result was consistent with previous studies of both wild and captive amphibians that reported dominance by one or few bacterial strains (Bates et al., 2018; Kueneman et al., 2016b, 2014; Loudon et al., 2014).

The Rubritaleaceae are a little studied family of Gram-negative bacteria in the phylum Verrucomicrobia, containing only five described species isolated from marine animals or marine sediment (Kasai et al., 2007; Rosenberg, 2014; Scheuermayer et al., 2006; Yoon et al., 2008, 2007). The 16S rRNA genes of these species are very highly conserved and the species are not distinguishable by 16S amplicon analysis (Rosenberg, 2014). This may explain why we were unable to assign taxonomy below the family level for the dominant Rubritaleaceae sequence variant on frogs. Described Rubritaleaceae species are non-motile, obligate aerobes that synthesize carotenoid pigments, resulting in red-colored colonies (Rosenberg, 2014). The production of carotenoid pigments by Rubritaleaceae on frog skin may affect the skin-associated microbial community, as previous studies have shown that dietary carotenoid intake by amphibians increased community richness and shifted community structure of frog skin microbiomes (Antwis et al., 2014; Edwards et al., 2017).

The only previous mention of the Rubritaleaceae in amphibian microbiomes was from a study of captive *Rana muscosa*, the sister species to *R. sierrae*, conducted in the same facility at the San Francisco Zoo as our study (Jani et al., 2021). However, the phylum Verrucomicrobia, which includes Rubritaleaceae, has been detected on amphibian skin in several studies based on 16S rRNA gene data (Becker et al., 2014; Belden et al., 2015; Kueneman et al., 2016b, 2014; Longo et al., 2015; Loudon et al., 2014; Sabino-Pinto et al., 2016; Sanchez et al., 2017). Verrucomicrobia is a widely distributed phylum that has been found in various environments including soil, marine and freshwater, and animal intestines (Bergmann et al., 2011; Hugenholtz et al., 1998; Parveen et al., 2013; Passel et al., 2011; van Passel et al., 2011). Although a previous study found that Verrucomicrobia were higher in relative abundance on wild than captive Panamanian golden frogs (*Atelopus zeteki*) (Becker et al., 2014), we hypothesize that the high relative abundance of Rubritaleaceae and Verrucomicrobia observed in the present study may be unique to captivity. This owes to the fact that Verrucomicrobia, though present, were not high in relative abundance in previous studies of wild *R. sierrae* populations (Jani and Briggs, 2014), even for populations from the same site used to source the San Francisco Zoo population examined here using the same primers for 16S rRNA gene amplification (Ellison et al., 2021, 2019). It is possible that in captivity, these taxa replace other taxa with similar functional abilities on the skin in the wild, but this requires further investigation.

The other dominant amphibian associated sequence variant was a member of the family Burkholderiaceae. This family consists of ecologically, phenotypically, and metabolically diverse Gram-negative bacteria found in soil, water, and in association with plants, animals, and fungi (Coenye, 2014). While some Burkholderiaceae are pathogens to plants and animals including humans (Coenye, 2014), others have been shown to suppress fungal pathogens (Carrión et al., 2018). The dominant Burkholderiaceae sequence variant observed here shared 100% sequence identity with a bacterial isolate from the *Bd* inhibitory database, suggesting that this bacterium may help suppress *Bd* proliferation on the skin (AmphiBac_1576 / Ranamuscosa-inhibitory_37; Woodhams et al., 2015).

The order Burkholderiales, which includes the family Burkholderiaceae, has been identified as highly relatively abundant on amphibians in several studies (Bataille et al., 2016; Bates et al., 2018; Kueneman et al., 2014), including studies of *R. sierrae* (Ellison et al., 2021, 2019). Previous research on wild *R. sierrae* found Burkholderiaceae on their skin, but it showed lower relative abundance compared to another family in the order, Comamonadaceae (Ellison et al., 2019). Notably, several amphibian microbiome studies report that a single Comamonadaceae sequence variant dominated the community in much the same way as the dominant Burkholderiaceae sequence variant did here (Bates et al., 2018; Kueneman et al., 2016b, 2014). Interestingly, while the taxonomic assignment for this bacterium was to Burkholderiaceae, the *Bd* inhibitory bacterial isolate with identical amplicon sequence was classified as an undescribed Comamonadaceae in the inhibitory database metadata (Woodhams et al., 2015). This discrepancy illustrates how choice of assignment algorithm and/or taxonomic database can impact such classifications. Therefore, the dominant sequence variant we identified as a Burkholderiaceae is likely to be closely related to the dominant Comamonadaceae found in other amphibian studies, or it may even represent the same bacterium. Despite amplicon sequence similarity to a known *Bd*-inhibitor, further work is needed to determine whether the specific taxon identified here exhibits anti-*Bd* function.

### 4.2 Tank identity was more strongly associated with skin microbiomes than frog individual identity

Next, we tested our hypothesis that frog individuals would vary in their microbiomes. However, because frog individuals were not distributed evenly among distinct tank environments, many comparisons of individuals were confounded with tank comparisons. This warranted an evaluation of the relative importance of individual and tank identity to observed variation. Therefore, we compared the proportion of comparisons that were significant within and between tanks for metrics of microbiome diversity, structure, and relative abundance of taxa.

We found that individuals housed in the same tanks did exhibit variation in all metrics. However, a greater percentage of between-tank comparisons were significantly different for all metrics except for relative Shannon diversity. This indicates that tank identity was more important than individual identity to variation in most metrics. Impacts of enclosure on animal microbiomes have been shown previously in other systems (Breen et al., 2019; Hildebrand et al., 2013). Further, amphibians have been shown to be impacted by their environment and evidence indicates that they select for rare environmental microbes on their skin (Loudon et al., 2014; Walke et al., 2014). Our results are consistent with these previous findings. For example, community structure of the microbiome varied among individuals within and between tanks, however, for all three metrics examined, the percent of significant between-tank comparisons was greater than the percent of significant within-tank comparisons (Figure 3D-F; *see Results*). Further, the percentage of between-tank individual comparisons that showed significant differences in relative abundances of the two dominant taxa on frogs (*i.e.,* the dominant Burkholderiaceae and Rubritaleaceae sequence variants) was roughly four times greater than the percentage of significant within-tank individual comparisons.

While tank identity appeared to be more important than individual to describing variation in many microbiome metrics, we did detect some effect of individual frog identity within tanks. Most published studies have not addressed differences in community composition among individuals, most likely because the majority of studies collected only one sample per individual. Generally, studies have focused on population or species-level variation, however variation between microbiomes of individuals may also be important to predicting disease dynamics. Individual variation could arise from factors such as diet, local habitat, microclimate, age, sex, or host genetics (Jiménez and Sommer, 2017). However, captive *R. sierrae* in our study had the same diet, were housed in highly similar same tank environments, were all the same age, and all originated from egg masses collected from the same population (while different egg masses could be associated with different genotypes, many frogs would have been siblings from the same egg mass). Additionally, we found that frog sex did not explain variation after controlling for differences between individuals. It is possible that the observed microbiome variation between individuals could be due to variation in efficacy of individual immune responses and in the amount or type of AMPs or other glandular secretions produced by individuals, and these differences require further study.

Considering that our study and Ellison et al (2021) both identify variation among *R. sierrae* individuals, we urge future researchers to collect replicate samples of individuals in order to document and identify sources of this variability, and to test whether individual variation in microbiomes and other components of the frog skin biome are important to within-population differences in *Bd*-driven disease outcomes. Further, future studies of microbiome variation among captive individuals should ensure that frogs are distributed among tank enclosures in a balanced design to more effectively tease apart the impacts of tank environment and individual identity.

### 4.3 Frog body regions showed spatial variation in the microbiome, which corresponded to expected spatial variation in *Bd* infection

Finally, we tested our hypotheses that we would detect differences in the microbiome between body regions of frogs, and further, that some variation would correspond to body regions preferentially infected by *Bd* despite the fact that these frogs were uninfected at the time of sampling (*see Supplementary Material*). We found evidence supporting these hypotheses in comparisons of within-sample diversity, community structure, and relative abundance of taxa, described below.

Within-sample diversity, for the most part, did not differ between frog body regions; the only exceptions were significantly higher relative Shannon diversity on frog forefeet compared to the abdomen, hindfeet, and back (Figure 2E, F). This differs from previous studies of the *Bombina orientalis* microbiome that found ventral surfaces harbored higher richness and diversity than dorsal surfaces (Bataille et al., 2016; Sabino-Pinto et al., 2016) but is consistent with findings from other amphibian species that showed no such differences (for *Bufo japonicus, Cynops pyrrhogaster, Odorrana splendida,* and *Rana japonica*) (Sabino-Pinto et al., 2016). Interestingly, there did not appear to be a relationship between the size of each body part and microbiome diversity. The forefeet were the smallest body region sampled, but harbored higher diversity than larger body regions like the back. This suggests that standardizing the number of strokes of each body region was sufficient to control for differences in the area occupied by each body region.

Microbial community structure differed primarily between the back and other body regions. Previous studies also found significant differences in microbiome structure across the skin for two different amphibian species (Bataille et al., 2016; Sabino-Pinto et al., 2016; Sanchez et al., 2017). Though in one study, community structure only differed significantly between dorsal and ventral surfaces in captivity and not in the wild, warranting future investigation of within-individual microbiome heterogeneity of wild *R. sierrae* (Bataille et al., 2016). Here, while we did not detect differences in unweighted community membership among body regions, we found that the back differed significantly from up to seven other body regions based on two metrics of relative abundance weighted community structure (Figure 3G-I). This suggests that shifts in dominant community members (*i.e,* those given more weight in these metrics) may be more important to differences between body regions than are differences in rare organisms or in presence/absence of organisms.

Supporting this claim, we also found that the relative abundance of the two dominant skin-associated bacteria (families Rubritaleaceae and Burkholderiaceae), both differed significantly in most comparisons of the back with other body regions (Figure 5B). As discussed above, the dominant Burkholderiaceae skin taxon was identified to be putatively *Bd*-inhibitory based on sequence similarity to a known anti-*Bd* bacterium (Woodhams et al., 2015). This Burkholderiaceae taxon had significantly higher relative abundance on body regions including the abdomen, inner hind-limbs, and hindfeet compared to the back. Further, we found that a different undescribed member of the Burkholderiaceae showed significant log2 fold higher normalized read counts on the abdomen compared to the back (SV56; Table S20), and this taxon was also putatively anti-*Bd* (Woodhams et al., 2015).

Our finding that much of what defines heterogeneity in the skin microbiome are differences between the back and other body regions supports our hypothesis that microbiome variation corresponds to spatial heterogeneity in *Bd* infection across the skin. Studies have shown that *Bd* infection occurs most on ventral surfaces, hindfeet, and toes, and is either absent or minimal (*i.e.,* very few *Bd* sporangia) on dorsal surfaces like the back (Berger et al., 2005, 1998; North and Alford, 2008; Pessier et al., 1999). Further, it has been shown that the back of an amphibian experiences fewer pathological changes due to *Bd* infection than do other body surfaces (Berger et al., 2005). The fact that two putatively anti-*Bd* members of the Burkolderiaceae showed higher relative abundance on ventral surfaces compared to the back suggests that they may directly interact with *Bd* upon infection. However, isolation and functional characterization of these taxa are needed to determine if they would act to inhibit *Bd* growth in practice. Additionally, as discussed above, our data is compositional, and therefore quantitative analyses of taxa are needed to determine whether differences in relative abundances of Burkholderiaceae observed were driven by differences in their absolute abundances or by differences in abundances of other taxa.

The reasons that both *Bd* infection and microbiome structure differ among body regions may relate to differences in skin architecture of the amphibian host. For example, there are usually larger and more numerous granular glands (also referred to as serous glands) on the back compared to ventral surfaces (Berger et al., 2005; Varga et al., 2019). Granular glands secrete bioactive molecules that assist in host defense, including AMPs (Varga et al., 2019). Such differences in the skin landscape may contribute to lower *Bd* infection on the back and to the differences in the skin microbiome between the back and other body regions that we observed here. Additionally, it has been shown that bacterially produced compounds can act synergistically with host-produced AMPs to inhibit *Bd* growth in *R. muscosa* (Myers et al., 2012). Thus, the combination of skin architecture and bacterial composition are likely directly relevant to the distribution of *Bd* infection across the skin.

Our results suggest that where you collect an amphibian skin swab from (*i.e.,* which body regions) will affect the resultant community observed. However, we emphasize that this may not apply to all types of amphibians. The heterogeneity in skin structure, microbiome structure, and *Bd* localization across the skin are all likely related to the evolved ecology of the amphibian. *R. sierrae* is a semi-aquatic species that spends much of its time basking at the edges of lakes and streams, keeping ventral surfaces and toes more moist than the back. *Bd* zoospores require water to disperse, so differences in moisture across the skin due to an amphibian’s lifestyle and ecology may contribute to spatial heterogeneity of *Bd* infection across the skin. For example, in a fully aquatic amphibian species (*Xenopus tropicalus*), no differences were detected in *Bd* infection between dorsal and ventral regions (Parker et al., 2002). Future studies would benefit from comparing differences in the microbiome structure, skin architecture, and spatial heterogeneity in *Bd* infection across amphibians of differing ecologies to determine whether there are consistent patterns and to elucidate the role that ecology has played in the evolution of such differences.

### 4.4 Applications for restoration

*R. sierrae* are critically endangered, but there are cases where management efforts have led to population recovery (Knapp et al., 2016). Knowledge about the microbiome could help us improve restoration efforts further (Redford et al., 2012). We suggest that microbiome variation between individuals, between distinct local environments, and within individuals could be important to restoration. Differences between individuals and between their local environments should be taken into consideration as they may lead to differences in outcomes after reintroduction and to differences in efficacy of probiotic treatments (*e.g.,* differences in successful colonization of the community using probiotic therapies). Differences across the skin could be exploited to focus on altering microbiomes of ventral surfaces and feet that gain higher *Bd* loads. Additionally, by elucidating microbiome variation between and within individuals, we can better understand and develop models to predict corresponding variation in *Bd* intensity. This natural variation relates to how susceptible frogs will be to high levels of infection (Ellison et al., 2019; Jani and Briggs, 2014), which could also be an indicator of how healthy and resilient frogs are in the face of other pathogens or environmental stressors.

## 5 Data Availability Statement

The raw 16S rRNA gene amplicon sequence data for this project has been deposited in the National Center for Biotechnology Information Sequence Read Archive under BioProject PRJNA1219149.

## 6 Author Contributions

SG and JE conceived of the study. SG designed the experiment, performed sampling, analyzed data, prepared figures and tables, and wrote and reviewed the drafts of the manuscript. JE advised on study design and data analyses and edited and reviewed the drafts of the manuscript.

## 9 Acknowledgements

We thank Jessie Bushell and the San Francisco Zoo for permitting and facilitating sampling. We thank Roland Knapp and the Mountain Lakes Research Group for assaying our swab samples for *Bd*. We thank Vance T. Vredenburg for providing advice on experimental design. We thank Cassandra L. Ettinger and John Jay Stachowicz for providing advice on analysis and on the manuscript. The work presented is derived from the doctoral dissertation of Sonia L. Ghose (Ghose, 2024).

