## Supplementary Material for "Skin microbiomes of frogs vary among individuals and body regions, revealing differences that reflect known patterns of chytrid infection"

### 1 Supplementary Methods and Results

#### 1.1 Assessing *Bd* infection

In addition to a microbiome swab sample, an additional swab sample was collected per individual to assess *Bd* presence and infection intensity (“*Bd* swabs”). To collect *Bd* swabs, we used a standard protocol, taking 30 strokes of ventral surfaces (10 strokes of ventral abdomen, 10 strokes of inner hindlimbs, and 5 strokes on each hindfoot; (“Chytrid Swabbing Protocol,” 2009).

DNA was extracted from *Bd* swab samples using the PrepMan Ultra Sample Preparation Reagent following the manufacturer’s protocol (40 μL used per sample). DNA extracts were stored at -20 °C or -80 °C. To confirm that frogs reared at the Zoo were not infected with *Bd*, we used a standard quantitative PCR (qPCR) assay with *Bd*-specific primers for ITS1 (Boyle et al., 2004; Hyatt et al., 2007). qPCR assays of DNA extracts were completed in Roland Knapp’s laboratory at the Sierra Nevada Aquatic Research Laboratory (Mammoth Lakes, California, USA). Plasmid *Bd* standards were used in assays, which are based on single ITS1 PCR amplicons (Joseph and Knapp, 2018; Longo et al., 2013).

No amplification of *Bd* ITS1 was detected during qPCR of frog *Bd* swab samples, indicating that *Bd* was not present on frogs sampled in this study.

### 2 Supplementary Figures

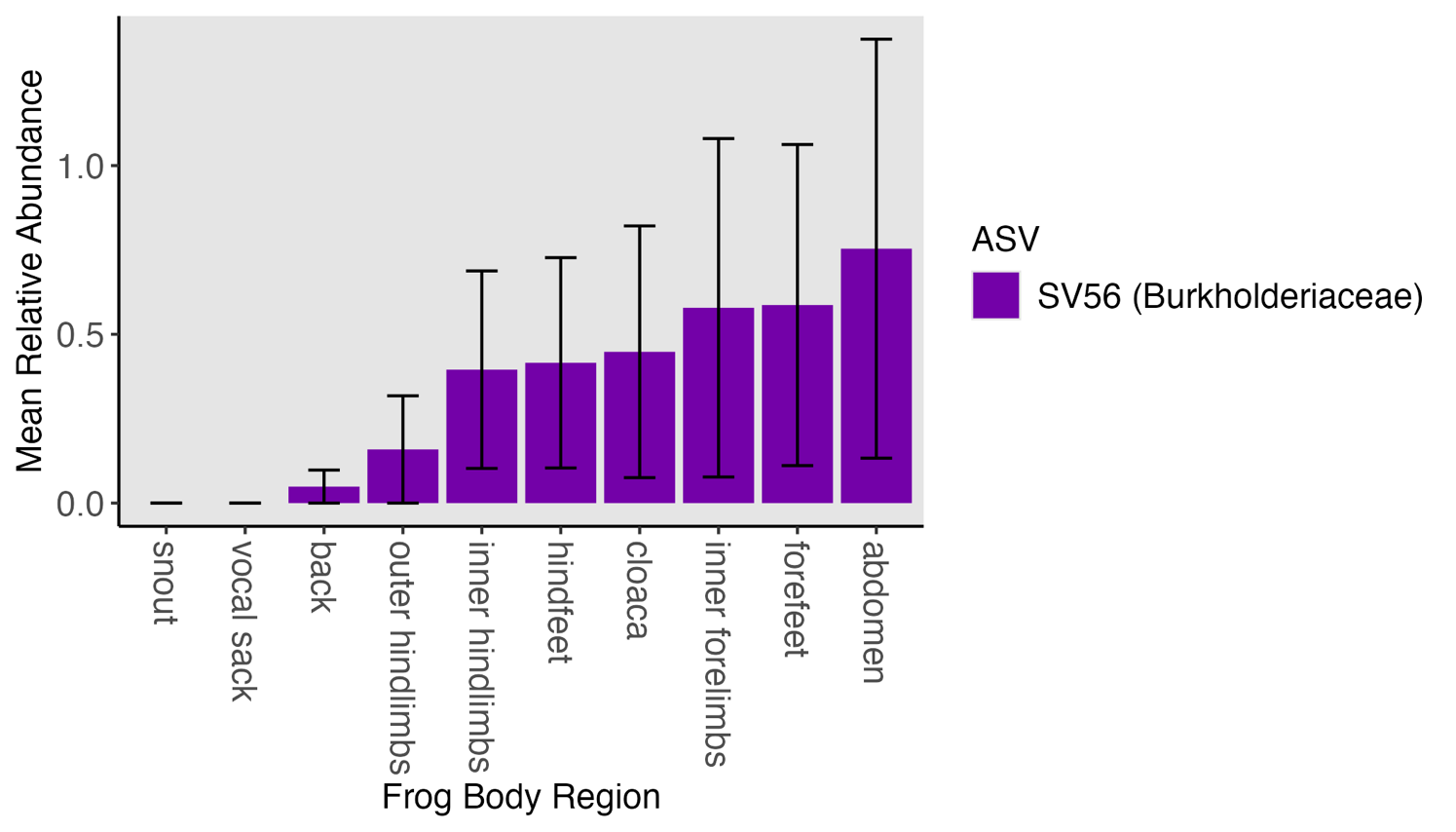

**Figure S1. Mean relative abundance of Burkholderiaceae taxon identified to have differential normalized counts between body regions.** SV56 had a significant log_2_ fold difference in DESeq2 normalized read counts between the abdomen and back (estimate of log_2_ fold difference = 24.18, p < 0.05; Table S20); Plot shows mean relative abundance of SV56 (family Burkholderiaceae) on frogs across all body regions calculated from the rarefied dataset; Frog body regions are ordered from lowest to highest mean relative abundance of SV56; Standard errors of means are represented with error bars; Mean relative abundance of SV56 was not significantly different between body regions (Kruskal Wallis, chi-squared = 5.5597, df = 9, p = 0.783).

### 3 Supplementary Tables

**Table S1. Pairwise comparisons of alpha diversity estimates among frog and environment sample types.** Following significant results of Kruskal-Wallis tests (Shannon: chi-squared = 56.771, df = 4, p < 0.001; Observed: chi-squared = 55.624, df = 4, p < 0.001), results of *post hoc* Dunn tests for Shannon diversity and observed richness in pairwise comparisons of frog, rock perch, tank wall, tank water, and underwater rock samples are shown; p-values were adjusted using Benjamini-Hochberg procedure to control the false discovery rate (FDR) with multiple comparisons; for significance, “***” = p ≤ 0.001, “**” = p ≤ 0.01, “*” = p ≤ 0.05, “ns” = not significant.

|  | **Comparison** | **Z** | **Unadjusted  P-value** | **Adjusted  P-value** | **Significance** |
| --- | --- | --- | --- | --- | --- |
| **Shannon** | frog ↔ rock perch | -3.851 | 0.00011771 | 0.00039238 | *** |
|  | frog ↔ tank wall | -4.966 | 0.00000068 | 0.00000682 | *** |
|  | frog ↔ tank water | -4.544 | 0.00000551 | 0.00002754 | *** |
|  | frog ↔ underwater rock | -2.569 | 0.01020177 | 0.02550442 | * |
|  | rock perch ↔ tank wall | -0.214 | 0.83080469 | 0.92311632 | ns |
|  | rock perch ↔ tank water | -0.507 | 0.61246706 | 1.00000000 | ns |
|  | rock perch ↔ underwater rock | 0.164 | 0.86972014 | 0.86972014 | ns |
|  | tank wall ↔ tank water | -0.341 | 0.73292772 | 0.91615965 | ns |
|  | tank wall ↔ underwater rock | 0.343 | 0.73168590 | 1.00000000 | ns |
|  | tank water ↔ underwater rock | 0.578 | 0.56352445 | 1.00000000 | ns |
| **Observed** | frog ↔ rock perch | -3.869 | 0.00010925 | 0.00036417 | *** |
|  | frog ↔ tank wall | -5.143 | 0.00000027 | 0.00000271 | *** |
|  | frog ↔ tank water | -4.216 | 0.00002485 | 0.00012425 | *** |
|  | frog ↔ underwater rock | -2.479 | 0.01318713 | 0.03296782 | * |
|  | rock perch ↔ tank wall | -0.316 | 0.75202298 | 1.00000000 | ns |
|  | rock perch ↔ tank water | -0.253 | 0.79989743 | 0.99987179 | ns |
|  | rock perch ↔ underwater rock | 0.250 | 0.80275295 | 0.89194772 | ns |
|  | tank wall ↔ tank water | 0.038 | 0.96944868 | 0.96944868 | ns |
|  | tank wall ↔ underwater rock | 0.515 | 0.60673714 | 1.00000000 | ns |
|  | tank water ↔ underwater rock | 0.457 | 0.64784982 | 1.00000000 | ns |

**Table S2. Pairwise comparisons of alpha diversity estimates among frog individuals.** Following significant results of Kruskal-Wallis tests (Shannon: chi-squared = 29.950, df = 8, p < 0.001; Observed: chi-squared = 40.841, df = 8, p < 0.001), results of *post hoc* Dunn tests for Shannon diversity and observed richness in pairwise comparisons of frog individuals are shown; p-values were adjusted using the Benjamini-Hochberg procedure to control the false discovery rate (FDR) with multiple comparisons; for significance, “***” = p ≤ 0.001, “**” = p ≤ 0.01, “*” = p ≤ 0.05, “ns” = not significant.

|  | **Comparison** | **Z** | **Unadjusted  P-value** | **Adjusted  P-value** | **Significance** |
| --- | --- | --- | --- | --- | --- |
| **Shannon** | Tahoe178 ↔ Tahoe181 | 3.193 | 0.00141005 | 0.01269048 | * |
|  | Tahoe159 ↔ Tahoe181 | 3.270 | 0.00107693 | 0.01292313 | * |
|  | Tahoe163 ↔ Tahoe181 | 3.535 | 0.00040785 | 0.01468248 | * |
|  | Tahoe159 ↔ Tahoe183 | 3.021 | 0.00251608 | 0.01811576 | * |
|  | Tahoe163 ↔ Tahoe183 | 3.287 | 0.00101356 | 0.01824401 | * |
|  | Tahoe178 ↔ Tahoe183 | 2.944 | 0.00323615 | 0.01941689 | * |
|  | Tahoe160 ↔ Tahoe163 | -2.850 | 0.00436894 | 0.02246885 | * |
|  | Tahoe123 ↔ Tahoe181 | 2.748 | 0.00600503 | 0.02702266 | * |
|  | Tahoe159 ↔ Tahoe160 | 2.585 | 0.00974125 | 0.03896500 | * |
|  | Tahoe123 ↔ Tahoe183 | 2.499 | 0.01244427 | 0.04072670 | * |
|  | Tahoe160 ↔ Tahoe178 | -2.508 | 0.01214687 | 0.04372875 | * |
|  | Tahoe172 ↔ Tahoe181 | 2.157 | 0.03101183 | 0.09303549 | ns |
|  | Tahoe162 ↔ Tahoe163 | -2.037 | 0.04164083 | 0.10707642 | ns |
|  | Tahoe123 ↔ Tahoe160 | 2.063 | 0.03913453 | 0.10837253 | ns |
|  | Tahoe172 ↔ Tahoe183 | 1.909 | 0.05630031 | 0.13512075 | ns |
|  | Tahoe159 ↔ Tahoe162 | 1.772 | 0.07643500 | 0.17197874 | ns |
|  | Tahoe162 ↔ Tahoe178 | -1.695 | 0.09012788 | 0.19085903 | ns |
|  | Tahoe160 ↔ Tahoe172 | -1.472 | 0.14097117 | 0.26710327 | ns |
|  | Tahoe162 ↔ Tahoe181 | 1.498 | 0.13416918 | 0.26833836 | ns |
|  | Tahoe163 ↔ Tahoe172 | 1.378 | 0.16819317 | 0.30274770 | ns |
|  | Tahoe123 ↔ Tahoe162 | 1.250 | 0.21142939 | 0.34597536 | ns |
|  | Tahoe162 ↔ Tahoe183 | 1.250 | 0.21142939 | 0.36245038 | ns |
|  | Tahoe159 ↔ Tahoe172 | 1.113 | 0.26583846 | 0.41609499 | ns |
|  | Tahoe172 ↔ Tahoe178 | -1.036 | 0.30035874 | 0.45053811 | ns |
|  | Tahoe123 ↔ Tahoe163 | -0.787 | 0.43102006 | 0.59679701 | ns |
|  | Tahoe160 ↔ Tahoe162 | -0.813 | 0.41614643 | 0.59925086 | ns |
|  | Tahoe162 ↔ Tahoe172 | -0.659 | 0.50985778 | 0.65553143 | ns |
|  | Tahoe160 ↔ Tahoe181 | 0.685 | 0.49351003 | 0.65801337 | ns |
|  | Tahoe123 ↔ Tahoe172 | 0.591 | 0.55479820 | 0.68871500 | ns |
|  | Tahoe123 ↔ Tahoe159 | -0.522 | 0.60159249 | 0.72191099 | ns |
|  | Tahoe160 ↔ Tahoe183 | 0.437 | 0.66245971 | 0.74526717 | ns |
|  | Tahoe123 ↔ Tahoe178 | -0.445 | 0.65626273 | 0.76211156 | ns |
|  | Tahoe163 ↔ Tahoe178 | 0.342 | 0.73207367 | 0.79862582 | ns |
|  | Tahoe181 ↔ Tahoe183 | -0.248 | 0.80396645 | 0.82693692 | ns |
|  | Tahoe159 ↔ Tahoe163 | -0.265 | 0.79075102 | 0.83726578 | ns |
|  | Tahoe159 ↔ Tahoe178 | 0.077 | 0.93859738 | 0.93859738 | ns |
| **Observed** | Tahoe163 ↔ Tahoe183 | 5.032 | 0.00000048 | 0.00001745 | *** |
|  | Tahoe163 ↔ Tahoe181 | 4.514 | 0.00000637 | 0.00011474 | *** |
|  | Tahoe160 ↔ Tahoe163 | -4.141 | 0.00003463 | 0.00041556 | *** |
|  | Tahoe123 ↔ Tahoe163 | -3.399 | 0.00067604 | 0.00608434 | ** |
|  | Tahoe172 ↔ Tahoe183 | 3.211 | 0.00132497 | 0.00953980 | ** |
|  | Tahoe162 ↔ Tahoe183 | 3.078 | 0.00208649 | 0.01073052 | * |
|  | Tahoe178 ↔ Tahoe183 | 3.116 | 0.00183189 | 0.01099134 | * |
|  | Tahoe172 ↔ Tahoe181 | 2.692 | 0.00710546 | 0.03197458 | * |
|  | Tahoe162 ↔ Tahoe181 | 2.559 | 0.01049791 | 0.03435680 | * |
|  | Tahoe159 ↔ Tahoe163 | -2.568 | 0.01024185 | 0.03687067 | * |
|  | Tahoe178 ↔ Tahoe181 | 2.598 | 0.00938888 | 0.03755550 | * |
|  | Tahoe159 ↔ Tahoe183 | 2.465 | 0.01371351 | 0.04114054 | * |
|  | Tahoe160 ↔ Tahoe172 | -2.319 | 0.02039809 | 0.05648702 | ns |
|  | Tahoe160 ↔ Tahoe178 | -2.225 | 0.02610523 | 0.06712772 | ns |
|  | Tahoe160 ↔ Tahoe162 | -2.186 | 0.02881084 | 0.06914600 | ns |
|  | Tahoe159 ↔ Tahoe181 | 1.946 | 0.05165154 | 0.10937972 | ns |
|  | Tahoe163 ↔ Tahoe178 | 1.916 | 0.05536231 | 0.11072463 | ns |
|  | Tahoe162 ↔ Tahoe163 | -1.955 | 0.05063036 | 0.11391832 | ns |
|  | Tahoe163 ↔ Tahoe172 | 1.822 | 0.06849733 | 0.12978441 | ns |
|  | Tahoe123 ↔ Tahoe183 | 1.633 | 0.10244391 | 0.18439903 | ns |
|  | Tahoe159 ↔ Tahoe160 | 1.573 | 0.11569343 | 0.18931652 | ns |
|  | Tahoe123 ↔ Tahoe172 | -1.577 | 0.11470442 | 0.19663615 | ns |
|  | Tahoe123 ↔ Tahoe178 | -1.483 | 0.13804911 | 0.21607687 | ns |
|  | Tahoe123 ↔ Tahoe162 | -1.445 | 0.14859343 | 0.22289015 | ns |
|  | Tahoe123 ↔ Tahoe181 | 1.114 | 0.26507992 | 0.38171508 | ns |
|  | Tahoe160 ↔ Tahoe183 | 0.892 | 0.37262255 | 0.51593891 | ns |
|  | Tahoe123 ↔ Tahoe159 | -0.832 | 0.40565619 | 0.54087491 | ns |
|  | Tahoe123 ↔ Tahoe160 | 0.742 | 0.45836148 | 0.56900045 | ns |
|  | Tahoe159 ↔ Tahoe172 | -0.746 | 0.45576769 | 0.58598703 | ns |
|  | Tahoe159 ↔ Tahoe178 | -0.652 | 0.51470238 | 0.61764285 | ns |
|  | Tahoe159 ↔ Tahoe162 | -0.613 | 0.53990574 | 0.62698731 | ns |
|  | Tahoe181 ↔ Tahoe183 | 0.519 | 0.60400157 | 0.67950176 | ns |
|  | Tahoe160 ↔ Tahoe181 | 0.373 | 0.70921021 | 0.77368387 | ns |
|  | Tahoe162 ↔ Tahoe172 | -0.133 | 0.89428949 | 0.94689476 | ns |
|  | Tahoe172 ↔ Tahoe178 | 0.094 | 0.92487017 | 0.95129504 | ns |
|  | Tahoe162 ↔ Tahoe178 | -0.039 | 0.96922714 | 0.96922714 | ns |

**Table S3. Pairwise comparisons of Shannon diversity among frog body regions.** Following significant results of Kruskal-Wallis test for differences in Shannon diversity (Shannon diversity: chi-squared = 21.543, df = 9, p = 0.01; Observed richness not significantly different: chi-squared = 8.674, df = 9, p = 0.468), results of *post hoc* Dunn tests for Shannon diversity in pairwise comparisons of frog body regions are shown; p-values were adjusted using Benjamini-Hochberg procedure to control the false discovery rate (FDR) with multiple comparisons; for significance, “***” = p ≤ 0.001, “**” = p ≤ 0.01, “*” = p ≤ 0.05, “ns” = not significant.

|  | **Comparison** | **Z** | **Unadjusted  P-value** | **Adjusted  P-value** | **Significance** |
| --- | --- | --- | --- | --- | --- |
| **Shannon** | abdomen ↔ forefeet | -3.762 | 0.00016839 | 0.00757741 | ** |
|  | forefeet ↔ hindfeet | 3.320 | 0.00089963 | 0.02024164 | * |
|  | back ↔ forefeet | -2.941 | 0.00326904 | 0.04903565 | * |
|  | forefeet ↔ inner hindlimbs | 2.589 | 0.00961514 | 0.10817030 | ns |
|  | abdomen ↔ snout | -2.382 | 0.01722542 | 0.12919061 | ns |
|  | abdomen ↔ vocal sack | -2.445 | 0.01448457 | 0.13036113 | ns |
|  | cloaca ↔ forefeet | -2.156 | 0.03105981 | 0.17471141 | ns |
|  | forefeet ↔ outer hindlimbs | 2.174 | 0.02967886 | 0.19079265 | ns |
|  | hindfeet ↔ vocal sack | -2.003 | 0.04518499 | 0.22592495 | ns |
|  | forefeet ↔ inner forelimbs | 1.877 | 0.06057054 | 0.22713953 | ns |
|  | hindfeet ↔ snout | -1.940 | 0.05240730 | 0.23583285 | ns |
|  | abdomen ↔ inner forelimbs | -1.886 | 0.05934353 | 0.24276898 | ns |
|  | back ↔ snout | -1.561 | 0.11856139 | 0.33345391 | ns |
|  | abdomen ↔ outer hindlimbs | -1.588 | 0.11230735 | 0.33692205 | ns |
|  | abdomen ↔ cloaca | -1.606 | 0.10828457 | 0.34805755 | ns |
|  | back ↔ vocal sack | -1.624 | 0.10437669 | 0.36130392 | ns |
|  | hindfeet ↔ inner forelimbs | -1.444 | 0.14886506 | 0.39405458 | ns |
|  | forefeet ↔ snout | 1.380 | 0.16746460 | 0.41866149 | ns |
|  | forefeet ↔ vocal sack | 1.317 | 0.18775782 | 0.44468957 | ns |
|  | inner hindlimbs ↔ vocal sack | -1.272 | 0.20332689 | 0.45748551 | ns |
|  | hindfeet ↔ outer hindlimbs | -1.146 | 0.25186992 | 0.47225610 | ns |
|  | cloaca ↔ hindfeet | 1.164 | 0.24447925 | 0.47832896 | ns |
|  | inner hindlimbs ↔ snout | -1.209 | 0.22667256 | 0.48572691 | ns |
|  | abdomen ↔ inner hindlimbs | -1.173 | 0.24084156 | 0.49263046 | ns |
|  | back ↔ inner forelimbs | -1.065 | 0.28704814 | 0.51668665 | ns |
|  | back ↔ outer hindlimbs | -0.767 | 0.44314874 | 0.62317791 | ns |
|  | inner forelimbs ↔ inner hindlimbs | 0.713 | 0.47599815 | 0.62999755 | ns |
|  | hindfeet ↔ inner hindlimbs | -0.731 | 0.46490246 | 0.63395789 | ns |
|  | cloaca ↔ snout | -0.776 | 0.43780263 | 0.63551995 | ns |
|  | back ↔ cloaca | -0.785 | 0.43249382 | 0.64874073 | ns |
|  | abdomen ↔ back | -0.821 | 0.41163480 | 0.66155592 | ns |
|  | outer hindlimbs ↔ snout | -0.794 | 0.42722248 | 0.66293143 | ns |
|  | cloaca ↔ vocal sack | -0.839 | 0.40143309 | 0.66905516 | ns |
|  | outer hindlimbs ↔ vocal sack | -0.857 | 0.39138468 | 0.67739656 | ns |
|  | inner forelimbs ↔ vocal sack | -0.559 | 0.57590489 | 0.74044915 | ns |
|  | inner forelimbs ↔ snout | -0.496 | 0.61973852 | 0.77467315 | ns |
|  | inner hindlimbs ↔ outer hindlimbs | -0.415 | 0.67812641 | 0.78245354 | ns |
|  | cloaca ↔ inner hindlimbs | 0.433 | 0.66496714 | 0.78746108 | ns |
|  | back ↔ hindfeet | 0.379 | 0.70473808 | 0.79283034 | ns |
|  | back ↔ inner hindlimbs | -0.352 | 0.72493893 | 0.79566468 | ns |
|  | abdomen ↔ hindfeet | -0.442 | 0.65842570 | 0.80078801 | ns |
|  | cloaca ↔ inner forelimbs | -0.280 | 0.77971675 | 0.81598264 | ns |
|  | inner forelimbs ↔ outer hindlimbs | 0.298 | 0.76590728 | 0.82061494 | ns |
|  | snout ↔ vocal sack | -0.063 | 0.94964276 | 0.97122555 | ns |
|  | cloaca ↔ outer hindlimbs | 0.018 | 0.98560343 | 0.98560343 | ns |

**Table S4. Variation in microbial community structure explained by frog and environment sample types.** Permutation multivariate analysis of variance (PERMANOVA) model outputs based on unweighted Unifrac distances, weighted Unifrac distances, and Bray-Curtis dissimilarities are shown; “EnvironType_FrogCategory” groupings include frog, rock perch, tank wall, tank water, and underwater rock samples.

|  | **Factor** | **Df** | **Sum of squares** | **Mean squares** | **F Model** | **R^2^** | **Pr(>F)** |
| --- | --- | --- | --- | --- | --- | --- | --- |
| **Unweighted Unifrac** | EnvironType_FrogCategory | 4 | 8.471 | 2.118 | 11.171 | 0.291 | 0.0001 |
|  | Residuals | 109 | 20.664 | 0.190 |  | 0.709 |  |
|  | Total | 113 | 29.136 |  |  | 1.000 |  |
| **Weighted Unifrac** | EnvironType_FrogCategory | 4 | 5.564 | 1.391 | 62.000 | 0.695 | 0.0001 |
|  | Residuals | 109 | 2.446 | 0.022 |  | 0.305 |  |
|  | Total | 113 | 8.009 |  |  | 1.000 |  |
| **Bray-Curtis** | EnvironType_FrogCategory | 4 | 10.214 | 2.553 | 22.456 | 0.452 | 0.0001 |
|  | Residuals | 109 | 12.394 | 0.114 |  | 0.548 |  |
|  | Total | 113 | 22.608 |  |  | 1.000 |  |

**Table S5. Pairwise comparisons of microbial community structure among frog and environment sample types.** Pairwise permutational multivariate analysis of variance (PERMANOVA) model outputs based on unweighted Unifrac distances, weighted Unifrac distances, and Bray-Curtis dissimilarities for frog, rock perch, tank wall, tank water, and underwater rock samples are shown; p-values were adjusted using the Benjamini-Hochberg procedure to control the false discovery rate (FDR) with multiple comparisons; “Signif.” indicates significance of adjusted p-values: “***” = p ≤ 0.001, “**” = p ≤ 0.01, “*” = p ≤ 0.05.

|  | **Combination** | **Sum of Squares** | **Mean Squares** | **F Model** | **R^2^** | **P-value** | **Adjust.  P-value** | **Signif** |
| --- | --- | --- | --- | --- | --- | --- | --- | --- |
| **Unweighted Unifrac** | frog ↔ rock perch | 2.101 | 2.101 | 10.850 | 0.103 | 0.0001 | 0.0003 | *** |
|  | frog ↔ tank wall | 3.470 | 3.470 | 18.187 | 0.158 | 0.0001 | 0.0003 | *** |
|  | frog ↔ tank water | 2.797 | 2.797 | 14.629 | 0.135 | 0.0001 | 0.0003 | *** |
|  | frog ↔ underwater rock | 1.191 | 1.191 | 6.149 | 0.063 | 0.0001 | 0.0003 | *** |
|  | rock perch ↔ tank wall | 0.514 | 0.514 | 2.936 | 0.184 | 0.0002 | 0.0004 | *** |
|  | rock perch ↔ tank water | 0.785 | 0.785 | 4.502 | 0.310 | 0.0025 | 0.0036 | ** |
|  | rock perch ↔ underwater rock | 0.412 | 0.412 | 2.061 | 0.227 | 0.0125 | 0.0125 | * |
|  | tank wall ↔ tank water | 0.659 | 0.659 | 4.179 | 0.243 | 0.0003 | 0.0005 | *** |
|  | tank wall ↔ underwater rock | 0.491 | 0.491 | 2.880 | 0.224 | 0.0053 | 0.0066 | ** |
|  | tank water ↔ underwater rock | 0.507 | 0.507 | 3.032 | 0.302 | 0.0118 | 0.0125 | * |
| **Weighted Unifrac** | frog ↔ rock perch | 1.678 | 1.678 | 80.771 | 0.462 | 0.0001 | 0.0003 | *** |
|  | frog ↔ tank wall | 1.791 | 1.791 | 95.433 | 0.496 | 0.0001 | 0.0003 | *** |
|  | frog ↔ tank water | 1.757 | 1.757 | 96.948 | 0.508 | 0.0001 | 0.0003 | *** |
|  | frog ↔ underwater rock | 1.093 | 1.093 | 62.537 | 0.407 | 0.0001 | 0.0003 | *** |
|  | rock perch ↔ tank wall | 0.399 | 0.399 | 7.493 | 0.366 | 0.0002 | 0.0004 | *** |
|  | rock perch ↔ tank water | 0.346 | 0.346 | 6.019 | 0.376 | 0.0025 | 0.0036 | ** |
|  | rock perch ↔ underwater rock | 0.176 | 0.176 | 2.664 | 0.276 | 0.0352 | 0.0352 | * |
|  | tank wall ↔ tank water | 0.411 | 0.411 | 12.082 | 0.482 | 0.0003 | 0.0005 | *** |
|  | tank wall ↔ underwater rock | 0.420 | 0.420 | 12.748 | 0.560 | 0.0053 | 0.0066 | ** |
|  | tank water ↔ underwater rock | 0.287 | 0.287 | 9.429 | 0.574 | 0.0118 | 0.0131 | * |
| **Bray-Curtis** | frog ↔ rock perch | 2.786 | 2.786 | 27.980 | 0.229 | 0.0001 | 0.0003 | *** |
|  | frog ↔ tank wall | 3.529 | 3.529 | 37.594 | 0.279 | 0.0001 | 0.0003 | *** |
|  | frog ↔ tank water | 3.123 | 3.123 | 33.940 | 0.265 | 0.0001 | 0.0003 | *** |
|  | frog ↔ underwater rock | 1.867 | 1.867 | 20.802 | 0.186 | 0.0001 | 0.0003 | *** |
|  | rock perch ↔ tank wall | 1.058 | 1.058 | 4.288 | 0.248 | 0.0002 | 0.0004 | *** |
|  | rock perch ↔ tank water | 1.132 | 1.132 | 4.116 | 0.292 | 0.0025 | 0.0036 | ** |
|  | rock perch ↔ underwater rock | 0.792 | 0.792 | 2.444 | 0.259 | 0.0125 | 0.0125 | * |
|  | tank wall ↔ tank water | 1.087 | 1.087 | 5.661 | 0.303 | 0.0003 | 0.0005 | *** |
|  | tank wall ↔ underwater rock | 1.080 | 1.080 | 5.364 | 0.349 | 0.0053 | 0.0066 | ** |
|  | tank water ↔ underwater rock | 0.922 | 0.922 | 4.141 | 0.372 | 0.0118 | 0.0125 | * |

**Table S6. Microbial community dispersion among frog and environmental sample types.** Results of permutation tests for homogeneity of multivariate dispersions based on unweighted Unifrac, weighted Unifrac, and Bray-Curtis dissimilarities for frog, rock perch, tank wall, tank water, and underwater rock samples (*i.e.,* “EnvironType_FrogCategory” groupings) are shown; “ns” = not significant, “*” = p ≤ 0.05, “**” = p ≤ 0.01**,** “***” = p ≤ 0.001.

|  | **Factor** | **Df** | **Sum of Squares** | **Mean Squares** | **F** | **No. Perm** | **Pr(>F)** |
| --- | --- | --- | --- | --- | --- | --- | --- |
| **Unweighted Unifrac Dispersion** | Groups (EnvironType_FrogCategory) | 4 | 0.056 | 0.014 | 1.709 | 9999 | 0.1583 ns |
|  | Residuals | 109 | 0.890 | 0.008 |  |  |  |
| **Weighted Unifrac Dispersion** | Groups (EnvironType_FrogCategory) | 4 | 0.136 | 0.034 | 12.755 | 9999 | 0.0001  *** |
|  | Residuals | 109 | 0.290 | 0.003 |  |  |  |
| **Bray Curtis Dispersion** | Groups (EnvironType_FrogCategory) | 4 | 0.638 | 0.159 | 12.443 | 9999 | 0.0001  *** |
|  | Residuals | 109 | 1.397 | 0.013 |  |  |  |

**Table S7. Pairwise comparisons of microbial community dispersion among frog and environmental sample types.** Results of *post hoc* Tukey multiple comparisons of mean dispersions based on weighted Unifrac and Bray-Curtis dissimilarities for frog, rock perch, tank wall, tank water, and underwater rock samples are shown; p-values were adjusted using Benjamini-Hochberg procedure to control the false discovery rate (FDR) with multiple comparisons; “Signif.” indicates significance of adjusted p-values: “ns” = not significant, “*” = p ≤ 0.05, “**” = p ≤ 0.01**,** “***” = p ≤ 0.001.

|  | **Combination** | **Difference** | **Lower 95% CI** | **Upper 95% CI** | **Adjusted  P-value** | **Signif.** |
| --- | --- | --- | --- | --- | --- | --- |
| **Weighted Unifrac Dispersion** | rock perch↔frog | 0.142 | 0.082 | 0.202 | 0.00000002 | *** |
|  | tank wall↔frog | 0.054 | 0.004 | 0.104 | 0.02957302 | * |
|  | tank water↔frog | 0.044 | -0.016 | 0.105 | 0.25749055 | ns |
|  | underwater rock↔frog | 0.008 | -0.076 | 0.091 | 0.99914303 | ns |
|  | tank wall↔rock perch | -0.089 | -0.164 | -0.013 | 0.01278466 | * |
|  | tank water↔rock perch | -0.098 | -0.180 | -0.015 | 0.01168291 | * |
|  | underwater rock↔rock perch | -0.135 | -0.236 | -0.033 | 0.00318957 | ** |
|  | tank water↔tank wall | -0.009 | -0.085 | 0.066 | 0.99697405 | ns |
|  | underwater rock↔tank wall | -0.046 | -0.141 | 0.049 | 0.66816610 | ns |
|  | underwater rock↔tank water | -0.037 | -0.138 | 0.064 | 0.85225254 | ns |
| **Bray Curtis Dispersion** | rock perch↔frog | 0.270 | 0.137 | 0.402 | 0.00000128 | *** |
|  | tank wall↔frog | 0.133 | 0.023 | 0.243 | 0.00949130 | ** |
|  | tank water↔frog | 0.145 | 0.012 | 0.277 | 0.02476239 | * |
|  | underwater rock↔frog | 0.149 | -0.035 | 0.333 | 0.17145704 | ns |
|  | tank wall↔rock perch | -0.137 | -0.302 | 0.029 | 0.15428667 | ns |
|  | tank water↔rock perch | -0.125 | -0.306 | 0.056 | 0.31732904 | ns |
|  | underwater rock↔rock perch | -0.121 | -0.343 | 0.101 | 0.56071676 | ns |
|  | tank water↔tank wall | 0.012 | -0.154 | 0.178 | 0.99963007 | ns |
|  | underwater rock↔tank wall | 0.016 | -0.193 | 0.226 | 0.99949948 | ns |
|  | underwater rock↔tank water | 0.004 | -0.218 | 0.226 | 0.99999793 | ns |

**Table S8. Variation in microbial community structure explained by frog individual and body region.** Permutational multivariate analysis of variance (PERMANOVA) model outputs based on unweighted Unifrac, weighted Unifrac, and Bray-Curtis dissimilarities for frog samples are shown; for significance, “ns” = not significant, “*” = p ≤ 0.05, “**” = p ≤ 0.01**,** “***” = p ≤ 0.001.

|  | **Factor** | **Df** | **Sums Of Squares** | **Mean Squares** | **F Model** | **R^2^** | **Pr(>F)** | **Significance** |
| --- | --- | --- | --- | --- | --- | --- | --- | --- |
| **Unweighted  Unifrac** | Body Region | 9 | 1.811 | 0.201 | 1.253 | 0.105 | 0.0440 | * |
|  | Individual | 8 | 3.844 | 0.481 | 2.993 | 0.223 | 0.0001 | *** |
|  | Residuals | 72 | 11.561 | 0.161 |  | 0.672 |  |  |
|  | Total | 89 | 17.216 |  |  | 1.000 |  |  |
| **Weighted  Unifrac** | Body Region | 9 | 0.431 | 0.048 | 7.091 | 0.279 | 0.0001 | *** |
|  | Individual | 8 | 0.625 | 0.078 | 11.574 | 0.405 | 0.0001 | *** |
|  | Residuals | 72 | 0.486 | 0.007 |  | 0.315 |  |  |
|  | Total | 89 | 1.541 |  |  | 1.000 |  |  |
| **Bray Curtis** | Body Region | 9 | 2.474 | 0.275 | 8.403 | 0.324 | 0.0001 | *** |
|  | Individual | 8 | 2.800 | 0.350 | 10.703 | 0.367 | 0.0001 | *** |
|  | Residuals | 72 | 2.355 | 0.033 |  | 0.309 |  |  |
|  | Total | 89 | 7.629 |  |  | 1.000 |  |  |

**Table S9. Pairwise comparisons of microbial community structure among frog individuals.** Pairwise permutational multivariate analysis of variance (PERMANOVA) model outputs based on unweighted Unifrac distances, weighted Unifrac distances, and Bray-Curtis dissimilarities for frog individuals are shown; p-values were adjusted using the Benjamini-Hochberg procedure to control the false discovery rate (FDR) with multiple comparisons; “Signif.” indicates significance of adjusted p-values: “***” = p ≤ 0.001, “**” = p ≤ 0.01, “*” = p ≤ 0.05, “ns” = not significant.

|  | **Combination** | **Sums of Squares** | **Mean Squares** | **F Model** | **R^2^** | **P-value** | **Adjust.  P-value** | **Signif.** |
| --- | --- | --- | --- | --- | --- | --- | --- | --- |
| **Unweighted Unifrac** | Tahoe123 ↔ Tahoe159 | 0.527 | 0.527 | 3.380 | 0.158 | 0.0008 | 0.0021 | ** |
|  | Tahoe123 ↔ Tahoe160 | 0.725 | 0.725 | 4.750 | 0.209 | 0.0001 | 0.0007 | *** |
|  | Tahoe123 ↔ Tahoe162 | 0.331 | 0.331 | 1.700 | 0.086 | 0.0694 | 0.0923 | ns |
|  | Tahoe123 ↔ Tahoe163 | 0.572 | 0.572 | 3.005 | 0.143 | 0.0002 | 0.0012 | ** |
|  | Tahoe123 ↔ Tahoe172 | 0.331 | 0.331 | 1.828 | 0.092 | 0.0109 | 0.0194 | * |
|  | Tahoe123 ↔ Tahoe178 | 0.540 | 0.540 | 3.379 | 0.158 | 0.0011 | 0.0025 | ** |
|  | Tahoe123 ↔ Tahoe181 | 0.440 | 0.440 | 2.890 | 0.138 | 0.0039 | 0.0074 | ** |
|  | Tahoe123 ↔ Tahoe183 | 0.194 | 0.194 | 1.201 | 0.063 | 0.2489 | 0.2658 | ns |
|  | Tahoe159 ↔ Tahoe160 | 0.169 | 0.169 | 1.249 | 0.065 | 0.2319 | 0.2658 | ns |
|  | Tahoe159 ↔ Tahoe162 | 0.642 | 0.642 | 3.609 | 0.167 | 0.0005 | 0.0015 | ** |
|  | Tahoe159 ↔ Tahoe163 | 0.750 | 0.750 | 4.328 | 0.194 | 0.0005 | 0.0015 | ** |
|  | Tahoe159 ↔ Tahoe172 | 0.317 | 0.317 | 1.936 | 0.097 | 0.0288 | 0.0415 | * |
|  | Tahoe159 ↔ Tahoe178 | 0.118 | 0.118 | 0.826 | 0.044 | 0.6171 | 0.6171 | ns |
|  | Tahoe159 ↔ Tahoe181 | 0.164 | 0.164 | 1.214 | 0.063 | 0.2510 | 0.2658 | ns |
|  | Tahoe159 ↔ Tahoe183 | 0.592 | 0.592 | 4.102 | 0.186 | 0.0001 | 0.0007 | *** |
|  | Tahoe160 ↔ Tahoe162 | 0.870 | 0.870 | 4.987 | 0.217 | 0.0001 | 0.0007 | *** |
|  | Tahoe160 ↔ Tahoe163 | 1.057 | 1.057 | 6.219 | 0.257 | 0.0003 | 0.0013 | ** |
|  | Tahoe160 ↔ Tahoe172 | 0.524 | 0.524 | 3.260 | 0.153 | 0.0014 | 0.0028 | ** |
|  | Tahoe160 ↔ Tahoe178 | 0.182 | 0.182 | 1.306 | 0.068 | 0.2224 | 0.2658 | ns |
|  | Tahoe160 ↔ Tahoe181 | 0.127 | 0.127 | 0.967 | 0.051 | 0.4380 | 0.4505 | ns |
|  | Tahoe160 ↔ Tahoe183 | 0.664 | 0.664 | 4.706 | 0.207 | 0.0006 | 0.0017 | ** |
|  | Tahoe162 ↔ Tahoe163 | 0.350 | 0.350 | 1.651 | 0.084 | 0.0718 | 0.0923 | ns |
|  | Tahoe162 ↔ Tahoe172 | 0.402 | 0.402 | 1.979 | 0.099 | 0.0307 | 0.0425 | * |
|  | Tahoe162 ↔ Tahoe178 | 0.601 | 0.601 | 3.308 | 0.155 | 0.0010 | 0.0024 | ** |
|  | Tahoe162 ↔ Tahoe181 | 0.630 | 0.630 | 3.616 | 0.167 | 0.0012 | 0.0025 | ** |
|  | Tahoe162 ↔ Tahoe183 | 0.498 | 0.498 | 2.714 | 0.131 | 0.0113 | 0.0194 | * |
|  | Tahoe163 ↔ Tahoe172 | 0.365 | 0.365 | 1.840 | 0.093 | 0.0197 | 0.0308 | * |
|  | Tahoe163 ↔ Tahoe178 | 0.685 | 0.685 | 3.863 | 0.177 | 0.0004 | 0.0015 | ** |
|  | Tahoe163 ↔ Tahoe181 | 0.864 | 0.864 | 5.094 | 0.221 | 0.0001 | 0.0007 | *** |
|  | Tahoe163 ↔ Tahoe183 | 0.865 | 0.865 | 4.836 | 0.212 | 0.0001 | 0.0007 | *** |
|  | Tahoe172 ↔ Tahoe178 | 0.261 | 0.261 | 1.553 | 0.079 | 0.0908 | 0.1127 | ns |
|  | Tahoe172 ↔ Tahoe181 | 0.350 | 0.350 | 2.186 | 0.108 | 0.0160 | 0.0262 | * |
|  | Tahoe172 ↔ Tahoe183 | 0.456 | 0.456 | 2.693 | 0.130 | 0.0005 | 0.0015 | ** |
|  | Tahoe178 ↔ Tahoe181 | 0.174 | 0.174 | 1.247 | 0.065 | 0.2400 | 0.2658 | ns |
|  | Tahoe178 ↔ Tahoe183 | 0.628 | 0.628 | 4.232 | 0.190 | 0.0003 | 0.0013 | ** |
|  | Tahoe181 ↔ Tahoe183 | 0.333 | 0.333 | 2.368 | 0.116 | 0.0218 | 0.0327 | * |
| **Weighted Unifrac** | Tahoe123 ↔ Tahoe159 | 0.016 | 0.016 | 1.573 | 0.080 | 0.2062 | 0.2395 | ns |
|  | Tahoe123 ↔ Tahoe160 | 0.045 | 0.045 | 3.685 | 0.170 | 0.0432 | 0.0676 | ns |
|  | Tahoe123 ↔ Tahoe162 | 0.090 | 0.090 | 10.802 | 0.375 | 0.0016 | 0.0058 | ** |
|  | Tahoe123 ↔ Tahoe163 | 0.229 | 0.229 | 13.470 | 0.428 | 0.0001 | 0.0012 | ** |
|  | Tahoe123 ↔ Tahoe172 | 0.020 | 0.020 | 2.466 | 0.120 | 0.0852 | 0.1136 | ns |
|  | Tahoe123 ↔ Tahoe178 | 0.108 | 0.108 | 9.588 | 0.348 | 0.0031 | 0.0093 | ** |
|  | Tahoe123 ↔ Tahoe181 | 0.010 | 0.010 | 1.658 | 0.084 | 0.1882 | 0.2258 | ns |
|  | Tahoe123 ↔ Tahoe183 | 0.063 | 0.063 | 6.405 | 0.262 | 0.0126 | 0.0267 | * |
|  | Tahoe159 ↔ Tahoe160 | 0.019 | 0.019 | 1.396 | 0.072 | 0.2424 | 0.2567 | ns |
|  | Tahoe159 ↔ Tahoe162 | 0.052 | 0.052 | 5.464 | 0.233 | 0.0234 | 0.0443 | * |
|  | Tahoe159 ↔ Tahoe163 | 0.121 | 0.121 | 6.663 | 0.270 | 0.0019 | 0.0062 | ** |
|  | Tahoe159 ↔ Tahoe172 | 0.043 | 0.043 | 4.741 | 0.208 | 0.0195 | 0.0390 | * |
|  | Tahoe159 ↔ Tahoe178 | 0.037 | 0.037 | 3.002 | 0.143 | 0.0638 | 0.0883 | ns |
|  | Tahoe159 ↔ Tahoe181 | 0.028 | 0.028 | 3.750 | 0.172 | 0.0419 | 0.0676 | ns |
|  | Tahoe159 ↔ Tahoe183 | 0.045 | 0.045 | 4.027 | 0.183 | 0.0418 | 0.0676 | ns |
|  | Tahoe160 ↔ Tahoe162 | 0.016 | 0.016 | 1.426 | 0.073 | 0.2301 | 0.2541 | ns |
|  | Tahoe160 ↔ Tahoe163 | 0.115 | 0.115 | 5.789 | 0.243 | 0.0053 | 0.0139 | * |
|  | Tahoe160 ↔ Tahoe172 | 0.086 | 0.086 | 7.859 | 0.304 | 0.0058 | 0.0139 | * |
|  | Tahoe160 ↔ Tahoe178 | 0.032 | 0.032 | 2.250 | 0.111 | 0.1250 | 0.1607 | ns |
|  | Tahoe160 ↔ Tahoe181 | 0.034 | 0.034 | 3.641 | 0.168 | 0.0546 | 0.0819 | ns |
|  | Tahoe160 ↔ Tahoe183 | 0.009 | 0.009 | 0.721 | 0.039 | 0.4172 | 0.4172 | ns |
|  | Tahoe162 ↔ Tahoe163 | 0.070 | 0.070 | 4.338 | 0.194 | 0.0056 | 0.0139 | * |
|  | Tahoe162 ↔ Tahoe172 | 0.163 | 0.163 | 23.319 | 0.564 | 0.0002 | 0.0018 | ** |
|  | Tahoe162 ↔ Tahoe178 | 0.018 | 0.018 | 1.715 | 0.087 | 0.1735 | 0.2154 | ns |
|  | Tahoe162 ↔ Tahoe181 | 0.084 | 0.084 | 15.810 | 0.468 | 0.0009 | 0.0041 | ** |
|  | Tahoe162 ↔ Tahoe183 | 0.008 | 0.008 | 0.844 | 0.045 | 0.3826 | 0.3935 | ns |
|  | Tahoe163 ↔ Tahoe172 | 0.303 | 0.303 | 19.364 | 0.518 | 0.0001 | 0.0012 | ** |
|  | Tahoe163 ↔ Tahoe178 | 0.027 | 0.027 | 1.428 | 0.073 | 0.2329 | 0.2541 | ns |
|  | Tahoe163 ↔ Tahoe181 | 0.270 | 0.270 | 19.292 | 0.517 | 0.0001 | 0.0012 | ** |
|  | Tahoe163 ↔ Tahoe183 | 0.111 | 0.111 | 6.293 | 0.259 | 0.0014 | 0.0056 | ** |
|  | Tahoe172 ↔ Tahoe178 | 0.164 | 0.164 | 16.478 | 0.478 | 0.0008 | 0.0041 | ** |
|  | Tahoe172 ↔ Tahoe181 | 0.021 | 0.021 | 4.314 | 0.193 | 0.0278 | 0.0500 | ns |
|  | Tahoe172 ↔ Tahoe183 | 0.129 | 0.129 | 15.101 | 0.456 | 0.0009 | 0.0041 | ** |
|  | Tahoe178 ↔ Tahoe181 | 0.134 | 0.134 | 16.137 | 0.473 | 0.0004 | 0.0029 | ** |
|  | Tahoe178 ↔ Tahoe183 | 0.036 | 0.036 | 3.051 | 0.145 | 0.0592 | 0.0852 | ns |
|  | Tahoe181 ↔ Tahoe183 | 0.056 | 0.056 | 8.117 | 0.311 | 0.0086 | 0.0194 | * |
| **Bray Curtis** | Tahoe123 ↔ Tahoe159 | 0.092 | 0.092 | 1.478 | 0.076 | 0.2220 | 0.2649 | ns |
|  | Tahoe123 ↔ Tahoe160 | 0.228 | 0.228 | 3.470 | 0.162 | 0.0570 | 0.0892 | ns |
|  | Tahoe123 ↔ Tahoe162 | 0.553 | 0.553 | 13.148 | 0.422 | 0.0009 | 0.0046 | ** |
|  | Tahoe123 ↔ Tahoe163 | 0.808 | 0.808 | 9.708 | 0.350 | 0.0002 | 0.0018 | ** |
|  | Tahoe123 ↔ Tahoe172 | 0.146 | 0.146 | 3.371 | 0.158 | 0.0436 | 0.0713 | ns |
|  | Tahoe123 ↔ Tahoe178 | 0.461 | 0.461 | 8.633 | 0.324 | 0.0042 | 0.0151 | * |
|  | Tahoe123 ↔ Tahoe181 | 0.040 | 0.040 | 1.154 | 0.060 | 0.2993 | 0.3265 | ns |
|  | Tahoe123 ↔ Tahoe183 | 0.373 | 0.373 | 7.246 | 0.287 | 0.0110 | 0.0283 | * |
|  | Tahoe159 ↔ Tahoe160 | 0.116 | 0.116 | 1.459 | 0.075 | 0.2281 | 0.2649 | ns |
|  | Tahoe159 ↔ Tahoe162 | 0.359 | 0.359 | 6.408 | 0.263 | 0.0147 | 0.0331 | * |
|  | Tahoe159 ↔ Tahoe163 | 0.395 | 0.395 | 4.064 | 0.184 | 0.0236 | 0.0472 | * |
|  | Tahoe159 ↔ Tahoe172 | 0.262 | 0.262 | 4.584 | 0.203 | 0.0269 | 0.0510 | ns |
|  | Tahoe159 ↔ Tahoe178 | 0.177 | 0.177 | 2.626 | 0.127 | 0.0838 | 0.1257 | ns |
|  | Tahoe159 ↔ Tahoe181 | 0.103 | 0.103 | 2.086 | 0.104 | 0.1414 | 0.1818 | ns |
|  | Tahoe159 ↔ Tahoe183 | 0.276 | 0.276 | 4.208 | 0.189 | 0.0427 | 0.0713 | ns |
|  | Tahoe160 ↔ Tahoe162 | 0.096 | 0.096 | 1.611 | 0.082 | 0.2048 | 0.2542 | ns |
|  | Tahoe160 ↔ Tahoe163 | 0.330 | 0.330 | 3.275 | 0.154 | 0.0417 | 0.0713 | ns |
|  | Tahoe160 ↔ Tahoe172 | 0.533 | 0.533 | 8.749 | 0.327 | 0.0052 | 0.0170 | * |
|  | Tahoe160 ↔ Tahoe178 | 0.099 | 0.099 | 1.389 | 0.072 | 0.2510 | 0.2824 | ns |
|  | Tahoe160 ↔ Tahoe181 | 0.121 | 0.121 | 2.291 | 0.113 | 0.1327 | 0.1769 | ns |
|  | Tahoe160 ↔ Tahoe183 | 0.056 | 0.056 | 0.812 | 0.043 | 0.3788 | 0.4011 | ns |
|  | Tahoe162 ↔ Tahoe163 | 0.284 | 0.284 | 3.681 | 0.170 | 0.0228 | 0.0472 | * |
|  | Tahoe162 ↔ Tahoe172 | 1.068 | 1.068 | 28.689 | 0.614 | 0.0001 | 0.0018 | ** |
|  | Tahoe162 ↔ Tahoe178 | 0.107 | 0.107 | 2.256 | 0.111 | 0.0924 | 0.1306 | ns |
|  | Tahoe162 ↔ Tahoe181 | 0.396 | 0.396 | 13.583 | 0.430 | 0.0019 | 0.0076 | ** |
|  | Tahoe162 ↔ Tahoe183 | 0.030 | 0.030 | 0.656 | 0.035 | 0.4333 | 0.4333 | ns |
|  | Tahoe163 ↔ Tahoe172 | 1.181 | 1.181 | 15.060 | 0.456 | 0.0002 | 0.0018 | ** |
|  | Tahoe163 ↔ Tahoe178 | 0.083 | 0.083 | 0.940 | 0.050 | 0.4007 | 0.4121 | ns |
|  | Tahoe163 ↔ Tahoe181 | 0.781 | 0.781 | 11.115 | 0.382 | 0.0002 | 0.0018 | ** |
|  | Tahoe163 ↔ Tahoe183 | 0.364 | 0.364 | 4.199 | 0.189 | 0.0109 | 0.0283 | * |
|  | Tahoe172 ↔ Tahoe178 | 0.819 | 0.819 | 16.847 | 0.483 | 0.0004 | 0.0029 | ** |
|  | Tahoe172 ↔ Tahoe181 | 0.189 | 0.189 | 6.255 | 0.258 | 0.0107 | 0.0283 | * |
|  | Tahoe172 ↔ Tahoe183 | 0.866 | 0.866 | 18.553 | 0.508 | 0.0008 | 0.0046 | ** |
|  | Tahoe178 ↔ Tahoe181 | 0.406 | 0.406 | 10.020 | 0.358 | 0.0011 | 0.0050 | ** |
|  | Tahoe178 ↔ Tahoe183 | 0.135 | 0.135 | 2.380 | 0.117 | 0.0943 | 0.1306 | ns |
|  | Tahoe181 ↔ Tahoe183 | 0.266 | 0.266 | 6.885 | 0.277 | 0.0145 | 0.0331 | * |

**Table S10. Microbial community dispersion by frog individual.** Results of permutation tests for homogeneity of multivariate dispersions based on unweighted Unifrac, weighted Unifrac, and Bray-Curtis dissimilarities for frog individuals are shown; for significance, “ns” = not significant, “*” = p ≤ 0.05, “**” = p ≤ 0.01**,** “***” = p ≤ 0.001.

|  | **Factor** | **Df** | **Sum of Squares** | **Mean Squares** | **F** | **No. Permutations** | **Pr(>F)** | **Significance** |
| --- | --- | --- | --- | --- | --- | --- | --- | --- |
| **Unweighted  Unifrac  Dispersion** | Individual | 8 | 0.119 | 0.015 | 2.198 | 9999 | 0.038 | * |
|  | Residuals | 81 | 0.548 | 0.007 |  |  |  |  |
| **Weighted  Unifrac  Dispersion** | Individual | 8 | 0.050 | 0.006 | 1.746 | 9999 | 0.096 | ns |
|  | Residuals | 81 | 0.290 | 0.004 |  |  |  |  |
| **Bray Curtis  Dispersion** | Individual | 8 | 0.246 | 0.031 | 1.572 | 9999 | 0.143 | ns |
|  | Residuals | 81 | 1.586 | 0.020 |  |  |  |  |

**Table S11. Pairwise comparisons of microbial community dispersion among frog individuals.** Results of *post hoc* Tukey multiple comparisons of mean dispersions based on unweighted Unifrac for frog individuals are shown; p-values were adjusted using Benjamini-Hochberg procedure to control the false discovery rate (FDR) with multiple comparisons; for significance, “ns” = not significant, “*” = p ≤ 0.05, “**” = p ≤ 0.01**,** “***” = p ≤ 0.001.

|  | **Combination** | **Difference** | **Lower 95% CI** | **Upper 95% CI** | **Adjusted P-value** | **Significance** |
| --- | --- | --- | --- | --- | --- | --- |
| **Unweighted Unifrac Dispersion** | Tahoe159 ↔ Tahoe123 | -0.050 | -0.167 | 0.067 | 0.911 | ns |
|  | Tahoe160 ↔ Tahoe123 | -0.057 | -0.174 | 0.061 | 0.834 | ns |
|  | Tahoe162 ↔ Tahoe123 | 0.045 | -0.072 | 0.163 | 0.946 | ns |
|  | Tahoe163 ↔ Tahoe123 | 0.031 | -0.087 | 0.148 | 0.996 | ns |
|  | Tahoe172 ↔ Tahoe123 | 0.011 | -0.106 | 0.128 | 1.000 | ns |
|  | Tahoe178 ↔ Tahoe123 | -0.037 | -0.154 | 0.080 | 0.984 | ns |
|  | Tahoe181 ↔ Tahoe123 | -0.055 | -0.172 | 0.063 | 0.860 | ns |
|  | Tahoe183 ↔ Tahoe123 | -0.032 | -0.149 | 0.085 | 0.994 | ns |
|  | Tahoe160 ↔ Tahoe159 | -0.007 | -0.124 | 0.111 | 1.000 | ns |
|  | Tahoe162 ↔ Tahoe159 | 0.095 | -0.022 | 0.213 | 0.207 | ns |
|  | Tahoe163 ↔ Tahoe159 | 0.080 | -0.037 | 0.198 | 0.425 | ns |
|  | Tahoe172 ↔ Tahoe159 | 0.061 | -0.057 | 0.178 | 0.774 | ns |
|  | Tahoe178 ↔ Tahoe159 | 0.013 | -0.104 | 0.130 | 1.000 | ns |
|  | Tahoe181 ↔ Tahoe159 | -0.005 | -0.122 | 0.113 | 1.000 | ns |
|  | Tahoe183 ↔ Tahoe159 | 0.018 | -0.100 | 0.135 | 1.000 | ns |
|  | Tahoe162 ↔ Tahoe160 | 0.102 | -0.015 | 0.219 | 0.140 | ns |
|  | Tahoe163 ↔ Tahoe160 | 0.087 | -0.030 | 0.204 | 0.315 | ns |
|  | Tahoe172 ↔ Tahoe160 | 0.067 | -0.050 | 0.185 | 0.660 | ns |
|  | Tahoe178 ↔ Tahoe160 | 0.020 | -0.098 | 0.137 | 1.000 | ns |
|  | Tahoe181 ↔ Tahoe160 | 0.002 | -0.115 | 0.119 | 1.000 | ns |
|  | Tahoe183 ↔ Tahoe160 | 0.025 | -0.093 | 0.142 | 0.999 | ns |
|  | Tahoe163 ↔ Tahoe162 | -0.015 | -0.132 | 0.102 | 1.000 | ns |
|  | Tahoe172 ↔ Tahoe162 | -0.035 | -0.152 | 0.083 | 0.990 | ns |
|  | Tahoe178 ↔ Tahoe162 | -0.082 | -0.200 | 0.035 | 0.390 | ns |
|  | Tahoe181 ↔ Tahoe162 | -0.100 | -0.217 | 0.017 | 0.158 | ns |
|  | Tahoe183 ↔ Tahoe162 | -0.078 | -0.195 | 0.040 | 0.476 | ns |
|  | Tahoe172 ↔ Tahoe163 | -0.020 | -0.137 | 0.098 | 1.000 | ns |
|  | Tahoe178 ↔ Tahoe163 | -0.068 | -0.185 | 0.050 | 0.658 | ns |
|  | Tahoe181 ↔ Tahoe163 | -0.085 | -0.202 | 0.032 | 0.345 | ns |
|  | Tahoe183 ↔ Tahoe163 | -0.063 | -0.180 | 0.055 | 0.743 | ns |
|  | Tahoe178 ↔ Tahoe172 | -0.048 | -0.165 | 0.069 | 0.928 | ns |
|  | Tahoe181 ↔ Tahoe172 | -0.065 | -0.183 | 0.052 | 0.695 | ns |
|  | Tahoe183 ↔ Tahoe172 | -0.043 | -0.160 | 0.074 | 0.961 | ns |
|  | Tahoe181 ↔ Tahoe178 | -0.018 | -0.135 | 0.100 | 1.000 | ns |
|  | Tahoe183 ↔ Tahoe178 | 0.005 | -0.112 | 0.122 | 1.000 | ns |
|  | Tahoe183 ↔ Tahoe181 | 0.023 | -0.095 | 0.140 | 0.999 | ns |

**Table S12. Pairwise comparisons of microbial community structure among frog body regions.** Pairwise permutational multivariate analysis of variance (PERMANOVA) model outputs based on unweighted Unifrac distances, weighted Unifrac distances, and Bray-Curtis dissimilarities for frog body regions are shown; p-values were adjusted using the Benjamini-Hochberg procedure to control the false discovery rate (FDR) with multiple comparisons; “Signif.” indicates significance of adjusted p-values: “***” = p ≤ 0.001, “**” = p ≤ 0.01, “*” = p ≤ 0.05, “ns” = not significant.

|  | **Combination** | **Sums of Squares** | **Mean Squares** | **F Model** | **R^2^** | **P-value** | **Adjust.  P-value** | **Signif.** |
| --- | --- | --- | --- | --- | --- | --- | --- | --- |
| **Unweighted  Unifrac** | abdomen ↔ back | 0.435 | 0.435 | 2.145 | 0.118 | 0.015 | 0.693 | ns |
|  | abdomen ↔ cloaca | 0.188 | 0.188 | 1.164 | 0.068 | 0.284 | 0.697 | ns |
|  | abdomen ↔ forefeet | 0.285 | 0.285 | 1.727 | 0.097 | 0.062 | 0.697 | ns |
|  | abdomen ↔ hindfeet | 0.204 | 0.204 | 1.297 | 0.075 | 0.217 | 0.697 | ns |
|  | abdomen ↔ inner forelimbs | 0.175 | 0.175 | 1.107 | 0.065 | 0.336 | 0.697 | ns |
|  | abdomen ↔ inner hindlimbs | 0.139 | 0.139 | 0.790 | 0.047 | 0.674 | 0.919 | ns |
|  | abdomen ↔ outer hindlimbs | 0.260 | 0.260 | 1.365 | 0.079 | 0.180 | 0.697 | ns |
|  | abdomen ↔ snout | 0.286 | 0.286 | 1.529 | 0.087 | 0.091 | 0.697 | ns |
|  | abdomen ↔ vocal sack | 0.259 | 0.259 | 1.511 | 0.086 | 0.098 | 0.697 | ns |
|  | back ↔ cloaca | 0.247 | 0.247 | 1.163 | 0.068 | 0.288 | 0.697 | ns |
|  | back ↔ forefeet | 0.235 | 0.235 | 1.087 | 0.064 | 0.334 | 0.697 | ns |
|  | back ↔ hindfeet | 0.231 | 0.231 | 1.108 | 0.065 | 0.317 | 0.697 | ns |
|  | back ↔ inner forelimbs | 0.279 | 0.279 | 1.334 | 0.077 | 0.185 | 0.697 | ns |
|  | back ↔ inner hindlimbs | 0.249 | 0.249 | 1.095 | 0.064 | 0.328 | 0.697 | ns |
|  | back ↔ outer hindlimbs | 0.265 | 0.265 | 1.097 | 0.064 | 0.337 | 0.697 | ns |
|  | back ↔ snout | 0.117 | 0.117 | 0.492 | 0.030 | 0.966 | 0.966 | ns |
|  | back ↔ vocal sack | 0.270 | 0.270 | 1.212 | 0.070 | 0.241 | 0.697 | ns |
|  | cloaca ↔ forefeet | 0.119 | 0.119 | 0.682 | 0.041 | 0.799 | 0.946 | ns |
|  | cloaca ↔ hindfeet | 0.113 | 0.113 | 0.678 | 0.041 | 0.772 | 0.946 | ns |
|  | cloaca ↔ inner forelimbs | 0.064 | 0.064 | 0.381 | 0.023 | 0.956 | 0.966 | ns |
|  | cloaca ↔ inner hindlimbs | 0.169 | 0.169 | 0.912 | 0.054 | 0.514 | 0.826 | ns |
|  | cloaca ↔ outer hindlimbs | 0.301 | 0.301 | 1.505 | 0.086 | 0.126 | 0.697 | ns |
|  | cloaca ↔ snout | 0.121 | 0.121 | 0.616 | 0.037 | 0.885 | 0.966 | ns |
|  | cloaca ↔ vocal sack | 0.140 | 0.140 | 0.774 | 0.046 | 0.694 | 0.919 | ns |
|  | forefeet ↔ hindfeet | 0.096 | 0.096 | 0.565 | 0.034 | 0.915 | 0.966 | ns |
|  | forefeet ↔ inner forelimbs | 0.140 | 0.140 | 0.815 | 0.048 | 0.649 | 0.912 | ns |
|  | forefeet ↔ inner hindlimbs | 0.225 | 0.225 | 1.186 | 0.069 | 0.273 | 0.697 | ns |
|  | forefeet ↔ outer hindlimbs | 0.379 | 0.379 | 1.860 | 0.104 | 0.052 | 0.697 | ns |
|  | forefeet ↔ snout | 0.110 | 0.110 | 0.551 | 0.033 | 0.942 | 0.966 | ns |
|  | forefeet ↔ vocal sack | 0.178 | 0.178 | 0.964 | 0.057 | 0.497 | 0.826 | ns |
|  | hindfeet ↔ inner forelimbs | 0.175 | 0.175 | 1.071 | 0.063 | 0.379 | 0.741 | ns |
|  | hindfeet ↔ inner hindlimbs | 0.151 | 0.151 | 0.828 | 0.049 | 0.590 | 0.912 | ns |
|  | hindfeet ↔ outer hindlimbs | 0.280 | 0.280 | 1.428 | 0.082 | 0.157 | 0.697 | ns |
|  | hindfeet ↔ snout | 0.098 | 0.098 | 0.510 | 0.031 | 0.955 | 0.966 | ns |
|  | hindfeet ↔ vocal sack | 0.146 | 0.146 | 0.829 | 0.049 | 0.635 | 0.912 | ns |
|  | inner forelimbs ↔ inner hindlimbs | 0.128 | 0.128 | 0.700 | 0.042 | 0.765 | 0.946 | ns |
|  | inner forelimbs ↔ outer hindlimbs | 0.304 | 0.304 | 1.542 | 0.088 | 0.122 | 0.697 | ns |
|  | inner forelimbs ↔ snout | 0.182 | 0.182 | 0.939 | 0.055 | 0.505 | 0.826 | ns |
|  | inner forelimbs ↔ vocal sack | 0.183 | 0.183 | 1.031 | 0.061 | 0.419 | 0.785 | ns |
|  | inner hindlimbs ↔ outer hindlimbs | 0.172 | 0.172 | 0.799 | 0.048 | 0.629 | 0.912 | ns |
|  | inner hindlimbs ↔ snout | 0.146 | 0.146 | 0.691 | 0.041 | 0.794 | 0.946 | ns |
|  | inner hindlimbs ↔ vocal sack | 0.191 | 0.191 | 0.973 | 0.057 | 0.484 | 0.826 | ns |
|  | outer hindlimbs ↔ snout | 0.268 | 0.268 | 1.186 | 0.069 | 0.265 | 0.697 | ns |
|  | outer hindlimbs ↔ vocal sack | 0.229 | 0.229 | 1.089 | 0.064 | 0.341 | 0.697 | ns |
|  | snout ↔ vocal sack | 0.123 | 0.123 | 0.596 | 0.036 | 0.934 | 0.966 | ns |
| **Weighted  Unifrac** | abdomen ↔ back | 0.117 | 0.117 | 8.998 | 0.360 | 0.003 | 0.019 | * |
|  | abdomen ↔ cloaca | 0.009 | 0.009 | 0.613 | 0.037 | 0.541 | 0.652 | ns |
|  | abdomen ↔ forefeet | 0.053 | 0.053 | 3.193 | 0.166 | 0.055 | 0.145 | ns |
|  | abdomen ↔ hindfeet | 0.008 | 0.008 | 0.595 | 0.036 | 0.462 | 0.630 | ns |
|  | abdomen ↔ inner forelimbs | 0.010 | 0.010 | 0.613 | 0.037 | 0.499 | 0.641 | ns |
|  | abdomen ↔ inner hindlimbs | 0.004 | 0.004 | 0.267 | 0.016 | 0.707 | 0.723 | ns |
|  | abdomen ↔ outer hindlimbs | 0.007 | 0.007 | 0.594 | 0.036 | 0.522 | 0.652 | ns |
|  | abdomen ↔ snout | 0.072 | 0.072 | 4.068 | 0.203 | 0.025 | 0.081 | ns |
|  | abdomen ↔ vocal sack | 0.026 | 0.026 | 1.433 | 0.082 | 0.229 | 0.368 | ns |
|  | back ↔ cloaca | 0.160 | 0.160 | 14.861 | 0.482 | 0.000 | 0.007 | ** |
|  | back ↔ forefeet | 0.177 | 0.177 | 13.637 | 0.460 | 0.001 | 0.008 | ** |
|  | back ↔ hindfeet | 0.184 | 0.184 | 19.896 | 0.554 | 0.000 | 0.007 | ** |
|  | back ↔ inner forelimbs | 0.103 | 0.103 | 8.172 | 0.338 | 0.002 | 0.015 | * |
|  | back ↔ inner hindlimbs | 0.138 | 0.138 | 12.648 | 0.441 | 0.001 | 0.011 | * |
|  | back ↔ outer hindlimbs | 0.106 | 0.106 | 12.131 | 0.431 | 0.001 | 0.011 | * |
|  | back ↔ snout | 0.022 | 0.022 | 1.581 | 0.090 | 0.172 | 0.322 | ns |
|  | back ↔ vocal sack | 0.064 | 0.064 | 4.407 | 0.216 | 0.012 | 0.055 | ns |
|  | cloaca ↔ forefeet | 0.023 | 0.023 | 1.568 | 0.089 | 0.222 | 0.368 | ns |
|  | cloaca ↔ hindfeet | 0.005 | 0.005 | 0.514 | 0.031 | 0.627 | 0.674 | ns |
|  | cloaca ↔ inner forelimbs | 0.010 | 0.010 | 0.682 | 0.041 | 0.492 | 0.641 | ns |
|  | cloaca ↔ inner hindlimbs | 0.002 | 0.002 | 0.197 | 0.012 | 0.882 | 0.882 | ns |
|  | cloaca ↔ outer hindlimbs | 0.009 | 0.009 | 0.891 | 0.053 | 0.413 | 0.599 | ns |
|  | cloaca ↔ snout | 0.085 | 0.085 | 5.558 | 0.258 | 0.008 | 0.040 | * |
|  | cloaca ↔ vocal sack | 0.030 | 0.030 | 1.884 | 0.105 | 0.155 | 0.302 | ns |
|  | forefeet ↔ hindfeet | 0.050 | 0.050 | 3.881 | 0.195 | 0.031 | 0.094 | ns |
|  | forefeet ↔ inner forelimbs | 0.022 | 0.022 | 1.357 | 0.078 | 0.256 | 0.397 | ns |
|  | forefeet ↔ inner hindlimbs | 0.032 | 0.032 | 2.229 | 0.122 | 0.117 | 0.264 | ns |
|  | forefeet ↔ outer hindlimbs | 0.036 | 0.036 | 2.920 | 0.154 | 0.067 | 0.167 | ns |
|  | forefeet ↔ snout | 0.081 | 0.081 | 4.615 | 0.224 | 0.022 | 0.075 | ns |
|  | forefeet ↔ vocal sack | 0.036 | 0.036 | 1.974 | 0.110 | 0.147 | 0.300 | ns |
|  | hindfeet ↔ inner forelimbs | 0.022 | 0.022 | 1.790 | 0.101 | 0.186 | 0.335 | ns |
|  | hindfeet ↔ inner hindlimbs | 0.006 | 0.006 | 0.515 | 0.031 | 0.550 | 0.652 | ns |
|  | hindfeet ↔ outer hindlimbs | 0.018 | 0.018 | 2.054 | 0.114 | 0.138 | 0.295 | ns |
|  | hindfeet ↔ snout | 0.117 | 0.117 | 8.477 | 0.346 | 0.002 | 0.015 | * |
|  | hindfeet ↔ vocal sack | 0.052 | 0.052 | 3.566 | 0.182 | 0.044 | 0.125 | ns |
|  | inner forelimbs ↔ inner hindlimbs | 0.006 | 0.006 | 0.453 | 0.028 | 0.591 | 0.665 | ns |
|  | inner forelimbs ↔ outer hindlimbs | 0.007 | 0.007 | 0.551 | 0.033 | 0.568 | 0.655 | ns |
|  | inner forelimbs ↔ snout | 0.044 | 0.044 | 2.582 | 0.139 | 0.093 | 0.220 | ns |
|  | inner forelimbs ↔ vocal sack | 0.007 | 0.007 | 0.394 | 0.024 | 0.666 | 0.697 | ns |
|  | inner hindlimbs ↔ outer hindlimbs | 0.005 | 0.005 | 0.461 | 0.028 | 0.629 | 0.674 | ns |
|  | inner hindlimbs ↔ snout | 0.077 | 0.077 | 4.980 | 0.237 | 0.016 | 0.065 | ns |
|  | inner hindlimbs ↔ vocal sack | 0.024 | 0.024 | 1.480 | 0.085 | 0.225 | 0.368 | ns |
|  | outer hindlimbs ↔ snout | 0.053 | 0.053 | 3.989 | 0.200 | 0.021 | 0.075 | ns |
|  | outer hindlimbs ↔ vocal sack | 0.016 | 0.016 | 1.150 | 0.067 | 0.310 | 0.464 | ns |
|  | snout ↔ vocal sack | 0.016 | 0.016 | 0.839 | 0.050 | 0.445 | 0.625 | ns |
| **Bray Curtis** | abdomen ↔ back | 0.682 | 0.682 | 10.461 | 0.395 | 0.004 | 0.017 | * |
|  | abdomen ↔ cloaca | 0.036 | 0.036 | 0.471 | 0.029 | 0.639 | 0.670 | ns |
|  | abdomen ↔ forefeet | 0.202 | 0.202 | 2.343 | 0.128 | 0.112 | 0.213 | ns |
|  | abdomen ↔ hindfeet | 0.049 | 0.049 | 0.697 | 0.042 | 0.424 | 0.564 | ns |
|  | abdomen ↔ inner forelimbs | 0.054 | 0.054 | 0.632 | 0.038 | 0.479 | 0.599 | ns |
|  | abdomen ↔ inner hindlimbs | 0.017 | 0.017 | 0.217 | 0.013 | 0.734 | 0.751 | ns |
|  | abdomen ↔ outer hindlimbs | 0.024 | 0.024 | 0.373 | 0.023 | 0.641 | 0.670 | ns |
|  | abdomen ↔ snout | 0.501 | 0.501 | 5.808 | 0.266 | 0.012 | 0.043 | * |
|  | abdomen ↔ vocal sack | 0.190 | 0.190 | 1.961 | 0.109 | 0.153 | 0.266 | ns |
|  | back ↔ cloaca | 0.867 | 0.867 | 19.843 | 0.554 | 0.000 | 0.004 | ** |
|  | back ↔ forefeet | 0.717 | 0.717 | 13.167 | 0.451 | 0.001 | 0.006 | ** |
|  | back ↔ hindfeet | 1.098 | 1.098 | 28.898 | 0.644 | 0.000 | 0.004 | ** |
|  | back ↔ inner forelimbs | 0.549 | 0.549 | 10.347 | 0.393 | 0.002 | 0.010 | ** |
|  | back ↔ inner hindlimbs | 0.794 | 0.794 | 16.407 | 0.506 | 0.001 | 0.006 | ** |
|  | back ↔ outer hindlimbs | 0.655 | 0.655 | 19.525 | 0.550 | 0.000 | 0.004 | ** |
|  | back ↔ snout | 0.075 | 0.075 | 1.383 | 0.080 | 0.244 | 0.379 | ns |
|  | back ↔ vocal sack | 0.269 | 0.269 | 4.139 | 0.206 | 0.019 | 0.056 | ns |
|  | cloaca ↔ forefeet | 0.103 | 0.103 | 1.590 | 0.090 | 0.203 | 0.338 | ns |
|  | cloaca ↔ hindfeet | 0.035 | 0.035 | 0.726 | 0.043 | 0.477 | 0.599 | ns |
|  | cloaca ↔ inner forelimbs | 0.043 | 0.043 | 0.675 | 0.040 | 0.520 | 0.616 | ns |
|  | cloaca ↔ inner hindlimbs | 0.015 | 0.015 | 0.249 | 0.015 | 0.860 | 0.860 | ns |
|  | cloaca ↔ outer hindlimbs | 0.029 | 0.029 | 0.668 | 0.040 | 0.569 | 0.640 | ns |
|  | cloaca ↔ snout | 0.584 | 0.584 | 9.010 | 0.360 | 0.000 | 0.004 | ** |
|  | cloaca ↔ vocal sack | 0.221 | 0.221 | 2.936 | 0.155 | 0.058 | 0.145 | ns |
|  | forefeet ↔ hindfeet | 0.269 | 0.269 | 4.553 | 0.222 | 0.025 | 0.067 | ns |
|  | forefeet ↔ inner forelimbs | 0.050 | 0.050 | 0.681 | 0.041 | 0.502 | 0.610 | ns |
|  | forefeet ↔ inner hindlimbs | 0.157 | 0.157 | 2.267 | 0.124 | 0.114 | 0.213 | ns |
|  | forefeet ↔ outer hindlimbs | 0.129 | 0.129 | 2.368 | 0.129 | 0.094 | 0.208 | ns |
|  | forefeet ↔ snout | 0.362 | 0.362 | 4.786 | 0.230 | 0.012 | 0.043 | * |
|  | forefeet ↔ vocal sack | 0.133 | 0.133 | 1.542 | 0.088 | 0.210 | 0.338 | ns |
|  | hindfeet ↔ inner forelimbs | 0.151 | 0.151 | 2.616 | 0.141 | 0.102 | 0.208 | ns |
|  | hindfeet ↔ inner hindlimbs | 0.042 | 0.042 | 0.784 | 0.047 | 0.400 | 0.563 | ns |
|  | hindfeet ↔ outer hindlimbs | 0.086 | 0.086 | 2.255 | 0.124 | 0.124 | 0.223 | ns |
|  | hindfeet ↔ snout | 0.836 | 0.836 | 14.151 | 0.469 | 0.000 | 0.004 | ** |
|  | hindfeet ↔ vocal sack | 0.389 | 0.389 | 5.597 | 0.259 | 0.016 | 0.051 | ns |
|  | inner forelimbs ↔ inner hindlimbs | 0.053 | 0.053 | 0.779 | 0.046 | 0.426 | 0.564 | ns |
|  | inner forelimbs ↔ outer hindlimbs | 0.031 | 0.031 | 0.575 | 0.035 | 0.555 | 0.640 | ns |
|  | inner forelimbs ↔ snout | 0.319 | 0.319 | 4.306 | 0.212 | 0.020 | 0.056 | ns |
|  | inner forelimbs ↔ vocal sack | 0.082 | 0.082 | 0.973 | 0.057 | 0.370 | 0.537 | ns |
|  | inner hindlimbs ↔ outer hindlimbs | 0.021 | 0.021 | 0.433 | 0.026 | 0.623 | 0.670 | ns |
|  | inner hindlimbs ↔ snout | 0.558 | 0.558 | 8.029 | 0.334 | 0.004 | 0.017 | * |
|  | inner hindlimbs ↔ vocal sack | 0.197 | 0.197 | 2.458 | 0.133 | 0.099 | 0.208 | ns |
|  | outer hindlimbs ↔ snout | 0.448 | 0.448 | 8.189 | 0.339 | 0.001 | 0.006 | ** |
|  | outer hindlimbs ↔ vocal sack | 0.146 | 0.146 | 2.234 | 0.123 | 0.102 | 0.208 | ns |
|  | snout ↔ vocal sack | 0.102 | 0.102 | 1.189 | 0.069 | 0.345 | 0.517 | ns |

**Table S13. Microbial community dispersion by frog body region.** Results of permutation tests for homogeneity of multivariate dispersions based on unweighted Unifrac, weighted Unifrac, and Bray-Curtis dissimilarities for frog body regions are shown; for significance, “ns” = not significant, “*” = p ≤ 0.05, “**” = p ≤ 0.01**,** “***” = p ≤ 0.001.

|  | **Factor** | **Df** | **Sum of Squares** | **Mean Squares** | **F** | **No. Permutations** | **Pr(>F)** | **Significance** |
| --- | --- | --- | --- | --- | --- | --- | --- | --- |
| **Unweighted Unifrac Dispersion** | Body Region | 9 | 0.110 | 0.012 | 1.369 | 9999 | 0.211 | ns |
|  | Residuals | 80 | 0.715 | 0.009 |  |  |  |  |
| **Weighted Unifrac Dispersion** | Body Region | 9 | 0.030 | 0.003 | 0.820 | 9999 | 0.605 | ns |
|  | Residuals | 80 | 0.321 | 0.004 |  |  |  |  |
| **Bray Curtis Dispersion** | Body Region | 9 | 0.185 | 0.021 | 1.191 | 9999 | 0.311 | ns |
|  | Residuals | 80 | 1.378 | 0.017 |  |  |  |  |

**Table S14. Pairwise comparisons of relative abundance of the family Burkholderiaceae among frog and environmental sample types.** Following significant results of Kruskal-Wallis test (Burkholderiaceae relative abundance: chi-squared = 357.31, df = 4, p < 0.001), results of *post hoc* Dunn tests for significant differences in relative abundance of the family Burkholderiaceae in pairwise comparisons of frog, rock perch, tank wall, tank water, and underwater rock samples are shown; p-values were adjusted using Benjamini-Hochberg procedure to control the false discovery rate (FDR) with multiple comparisons; for significance, “***” = p ≤ 0.001, “**” = p ≤ 0.01, “*” = p ≤ 0.05, “ns” = not significant.

| **Comparison** | **Z** | **Unadjusted  P-value** | **Adjusted  P-value** | **Significance** |
| --- | --- | --- | --- | --- |
| frog ↔ rock perch | -14.531 | 7.73E-48 | 7.73E-47 | *** |
| frog ↔ tank wall | -7.349 | 1.99E-13 | 6.65E-13 | *** |
| rock perch ↔ tank wall | 6.750 | 1.48E-11 | 2.96E-11 | *** |
| frog ↔ tank water | -9.939 | 2.83E-23 | 1.41E-22 | *** |
| rock perch ↔ tank water | 3.354 | 7.98E-04 | 1.33E-03 | ** |
| tank wall ↔ tank water | -3.076 | 2.10E-03 | 3.00E-03 | ** |
| frog ↔ underwater rock | -6.984 | 2.87E-12 | 7.18E-12 | *** |
| rock perch ↔ underwater rock | 2.868 | 4.13E-03 | 5.16E-03 | ** |
| tank wall ↔ underwater rock | -2.294 | 2.18E-02 | 2.42E-02 | * |
| tank water ↔ underwater rock | 0.130 | 8.97E-01 | 8.97E-01 | ns |

**Table S15. Pairwise comparisons of relative abundances of amplicon sequence variants among frog and environmental sample types.** Following significant results of Kruskal-Wallis tests (SV2 relative abundance: chi-squared = 56.492, df = 4, p < 0.001; SV1 relative abundance: chi-squared = 45.626, df = 4, p < 0.001), results of *post hoc* Dunn tests for significant differences in relative abundance of SV1 (family Rubritaleaceae) and SV2 (family Burkholderiaceae) in pairwise comparisons of frog, rock perch, tank wall, tank water, and underwater rock samples are shown; p-values were adjusted using Benjamini-Hochberg procedure to control the false discovery rate (FDR) with multiple comparisons; for significance, “***” = p ≤ 0.001, “**” = p ≤ 0.01, “*” = p ≤ 0.05, “ns” = not significant.

|  | **Comparison** | **Z** | **Unadjusted P-value** | **Adjusted P-value** | **Significance** |
| --- | --- | --- | --- | --- | --- |
| **SV2** | frog ↔ rock perch | 4.335 | 0.000015 | 0.000073 | *** |
|  | frog ↔ tank wall | 4.736 | 0.000002 | 0.000022 | *** |
|  | rock perch ↔ tank wall | -0.327 | 0.743755 | 1.000000 | ns |
|  | frog ↔ tank water | 3.982 | 0.000068 | 0.000227 | *** |
|  | rock perch ↔ tank water | -0.258 | 0.796680 | 1.000000 | ns |
|  | tank wall ↔ tank water | 0.045 | 0.964388 | 1.000000 | ns |
|  | frog ↔ underwater rock | 3.093 | 0.001981 | 0.004953 | ** |
|  | rock perch ↔ underwater rock | -0.018 | 0.985776 | 0.985776 | ns |
|  | tank wall ↔ underwater rock | 0.240 | 0.810706 | 1.000000 | ns |
|  | tank water ↔ underwater rock | 0.193 | 0.847320 | 1.000000 | ns |
| **SV1** | frog ↔ rock perch | 4.233 | 0.000023 | 0.000230 | *** |
|  | frog ↔ tank wall | 3.459 | 0.000543 | 0.001809 | ** |
|  | rock perch ↔ tank wall | -1.092 | 0.274712 | 0.457854 | ns |
|  | frog ↔ tank water | 3.575 | 0.000350 | 0.001748 | ** |
|  | rock perch ↔ tank water | -0.480 | 0.630973 | 0.701081 | ns |
|  | tank wall ↔ tank water | 0.566 | 0.571346 | 0.714183 | ns |
|  | frog ↔ underwater rock | 3.320 | 0.000899 | 0.002247 | ** |
|  | rock perch ↔ underwater rock | 0.232 | 0.816724 | 0.816724 | ns |
|  | tank wall ↔ underwater rock | 1.109 | 0.267284 | 0.534568 | ns |
|  | tank water ↔ underwater rock | 0.624 | 0.532647 | 0.760924 | ns |

**Table S16. Pairwise comparisons of relative abundances of amplicon sequence variants among frog individuals.** Following significant results of Kruskal-Wallis test (SV2 relative abundance: chi-squared = 27.863, df = 8, p < 0.001; SV1 relative abundance: chi-squared = 40.828, df = 8, p < 0.001), results of *post hoc* Dunn tests for significant differences in relative abundance of SV1 (Rubritaleaceae) and SV2 (Burkholderiaceae) in pairwise comparisons of frog individuals are shown; p-values were adjusted using Benjamini-Hochberg procedure to control the false discovery rate (FDR) with multiple comparisons; for significance, “***” = p ≤ 0.001, “**” = p ≤ 0.01, “*” = p ≤ 0.05, “ns” = not significant.

|  | **Comparison** | **Z** | **Unadjusted  P-value** | **Adjusted  P-value** | **Significance** |
| --- | --- | --- | --- | --- | --- |
| **SV2** | Tahoe123 ↔ Tahoe159 | -0.629 | 0.5292789 | 0.6146465 | ns |
|  | Tahoe123 ↔ Tahoe160 | -1.823 | 0.0682842 | 0.1638821 | ns |
|  | Tahoe159 ↔ Tahoe160 | -1.194 | 0.2324703 | 0.4184465 | ns |
|  | Tahoe123 ↔ Tahoe162 | -2.979 | 0.0028954 | 0.0208469 | * |
|  | Tahoe159 ↔ Tahoe162 | -2.350 | 0.0187975 | 0.0845888 | ns |
|  | Tahoe160 ↔ Tahoe162 | -1.156 | 0.2478846 | 0.4249450 | ns |
|  | Tahoe123 ↔ Tahoe163 | -1.682 | 0.0925885 | 0.2083241 | ns |
|  | Tahoe159 ↔ Tahoe163 | -1.053 | 0.2924366 | 0.4211087 | ns |
|  | Tahoe160 ↔ Tahoe163 | 0.141 | 0.8876897 | 0.9399068 | ns |
|  | Tahoe162 ↔ Tahoe163 | 1.297 | 0.1947238 | 0.4123562 | ns |
|  | Tahoe123 ↔ Tahoe172 | 1.220 | 0.2225796 | 0.4451592 | ns |
|  | Tahoe159 ↔ Tahoe172 | 1.849 | 0.0644860 | 0.1934581 | ns |
|  | Tahoe160 ↔ Tahoe172 | 3.043 | 0.0023437 | 0.0210932 | * |
|  | Tahoe162 ↔ Tahoe172 | 4.198 | 0.0000269 | 0.0009680 | *** |
|  | Tahoe163 ↔ Tahoe172 | 2.902 | 0.0037127 | 0.0222760 | * |
|  | Tahoe123 ↔ Tahoe178 | -1.836 | 0.0663627 | 0.1706470 | ns |
|  | Tahoe159 ↔ Tahoe178 | -1.207 | 0.2274866 | 0.4310273 | ns |
|  | Tahoe160 ↔ Tahoe178 | -0.013 | 0.9897563 | 0.9897563 | ns |
|  | Tahoe162 ↔ Tahoe178 | 1.143 | 0.2531782 | 0.4142915 | ns |
|  | Tahoe163 ↔ Tahoe178 | -0.154 | 0.8775569 | 0.9573348 | ns |
|  | Tahoe172 ↔ Tahoe178 | -3.056 | 0.0022456 | 0.0269476 | * |
|  | Tahoe123 ↔ Tahoe181 | -0.740 | 0.4590708 | 0.5508850 | ns |
|  | Tahoe159 ↔ Tahoe181 | -0.111 | 0.9114017 | 0.9374418 | ns |
|  | Tahoe160 ↔ Tahoe181 | 1.083 | 0.2789199 | 0.4183798 | ns |
|  | Tahoe162 ↔ Tahoe181 | 2.238 | 0.0252046 | 0.1008186 | ns |
|  | Tahoe163 ↔ Tahoe181 | 0.942 | 0.3464380 | 0.4796834 | ns |
|  | Tahoe172 ↔ Tahoe181 | -1.960 | 0.0499870 | 0.1799532 | ns |
|  | Tahoe178 ↔ Tahoe181 | 1.096 | 0.2732592 | 0.4277100 | ns |
|  | Tahoe123 ↔ Tahoe183 | -2.585 | 0.0097406 | 0.0500948 | ns |
|  | Tahoe159 ↔ Tahoe183 | -1.956 | 0.0504892 | 0.1652375 | ns |
|  | Tahoe160 ↔ Tahoe183 | -0.762 | 0.4461937 | 0.5736777 | ns |
|  | Tahoe162 ↔ Tahoe183 | 0.394 | 0.6937827 | 0.7805055 | ns |
|  | Tahoe163 ↔ Tahoe183 | -0.903 | 0.3665237 | 0.4886983 | ns |
|  | Tahoe172 ↔ Tahoe183 | -3.805 | 0.0001420 | 0.0025566 | ** |
|  | Tahoe178 ↔ Tahoe183 | -0.749 | 0.4538952 | 0.5634561 | ns |
|  | Tahoe181 ↔ Tahoe183 | -1.845 | 0.0651067 | 0.1802954 | ns |
| **SV1** | Tahoe123 ↔ Tahoe159 | 0.907 | 0.3642548 | 0.4521784 | ns |
|  | Tahoe123 ↔ Tahoe160 | 1.451 | 0.1468345 | 0.2298278 | ns |
|  | Tahoe159 ↔ Tahoe160 | 0.544 | 0.5867742 | 0.6401173 | ns |
|  | Tahoe123 ↔ Tahoe162 | 2.491 | 0.0127470 | 0.0382410 | * |
|  | Tahoe159 ↔ Tahoe162 | 1.583 | 0.1133136 | 0.2039645 | ns |
|  | Tahoe160 ↔ Tahoe162 | 1.040 | 0.2983600 | 0.4131138 | ns |
|  | Tahoe123 ↔ Tahoe163 | 4.066 | 0.0000479 | 0.0005747 | *** |
|  | Tahoe159 ↔ Tahoe163 | 3.158 | 0.0015864 | 0.0095187 | ** |
|  | Tahoe160 ↔ Tahoe163 | 2.615 | 0.0089261 | 0.0321340 | * |
|  | Tahoe162 ↔ Tahoe163 | 1.575 | 0.1152763 | 0.1976165 | ns |
|  | Tahoe123 ↔ Tahoe172 | -0.942 | 0.3464360 | 0.4454178 | ns |
|  | Tahoe159 ↔ Tahoe172 | -1.849 | 0.0644849 | 0.1365563 | ns |
|  | Tahoe160 ↔ Tahoe172 | -2.392 | 0.0167418 | 0.0463620 | * |
|  | Tahoe162 ↔ Tahoe172 | -3.432 | 0.0005985 | 0.0043093 | ** |
|  | Tahoe163 ↔ Tahoe172 | -5.007 | 0.0000006 | 0.0000199 | *** |
|  | Tahoe123 ↔ Tahoe178 | 2.615 | 0.0089261 | 0.0292128 | * |
|  | Tahoe159 ↔ Tahoe178 | 1.708 | 0.0877137 | 0.1661943 | ns |
|  | Tahoe160 ↔ Tahoe178 | 1.164 | 0.2443969 | 0.3519315 | ns |
|  | Tahoe162 ↔ Tahoe178 | 0.124 | 0.9012281 | 0.9012281 | ns |
|  | Tahoe163 ↔ Tahoe178 | -1.451 | 0.1468345 | 0.2402746 | ns |
|  | Tahoe172 ↔ Tahoe178 | 3.556 | 0.0003760 | 0.0033838 | ** |
|  | Tahoe123 ↔ Tahoe181 | -0.265 | 0.7907485 | 0.8133413 | ns |
|  | Tahoe159 ↔ Tahoe181 | -1.173 | 0.2409457 | 0.3614186 | ns |
|  | Tahoe160 ↔ Tahoe181 | -1.716 | 0.0861359 | 0.1722718 | ns |
|  | Tahoe162 ↔ Tahoe181 | -2.756 | 0.0058495 | 0.0233981 | * |
|  | Tahoe163 ↔ Tahoe181 | -4.331 | 0.0000148 | 0.0002672 | *** |
|  | Tahoe172 ↔ Tahoe181 | 0.676 | 0.4989226 | 0.5793940 | ns |
|  | Tahoe178 ↔ Tahoe181 | -2.880 | 0.0039741 | 0.0204383 | * |
|  | Tahoe123 ↔ Tahoe183 | 1.849 | 0.0644849 | 0.1450911 | ns |
|  | Tahoe159 ↔ Tahoe183 | 0.942 | 0.3464360 | 0.4619147 | ns |
|  | Tahoe160 ↔ Tahoe183 | 0.398 | 0.6906242 | 0.7312491 | ns |
|  | Tahoe162 ↔ Tahoe183 | -0.642 | 0.5209065 | 0.5860198 | ns |
|  | Tahoe163 ↔ Tahoe183 | -2.217 | 0.0266325 | 0.0684835 | ns |
|  | Tahoe172 ↔ Tahoe183 | 2.790 | 0.0052653 | 0.0236940 | * |
|  | Tahoe178 ↔ Tahoe183 | -0.766 | 0.4436414 | 0.5323696 | ns |
|  | Tahoe181 ↔ Tahoe183 | 2.114 | 0.0345024 | 0.0828057 | ns |

**Table S17. Comparison of SV1 and SV2 relative abundance within each frog individual.** Results of Wilcoxon signed rank tests for significant differences between relative abundances of SV1 (family Rubritaleaceae) and SV2 (family Burkholderiaceae) by frog individual; p-values were adjusted using the Benjamini-Hochberg procedure to control the false discovery rate (FDR) with multiple comparisons; for significance, “***” = p ≤ 0.001, “**” = p ≤ 0.01, “*” = p ≤ 0.05, “ns” = not significant.

| **Individual** | **Vstats** | **Estimate** | **Lower 95% CI** | **Upper 95% CI** | **P-value** | **Adjusted  P-value** | **Significance** |
| --- | --- | --- | --- | --- | --- | --- | --- |
| Tahoe172 | 2 | -44.609 | -64.081 | -20.389 | 0.0059 | 0.0410 | * |
| Tahoe181 | 14 | -12.486 | -34.855 | 7.572 | 0.1934 | 0.2486 | ns |
| Tahoe123 | 11 | -21.250 | -42.153 | 2.512 | 0.1055 | 0.1582 | ns |
| Tahoe159 | 25 | -4.254 | -36.450 | 31.170 | 0.8457 | 0.8457 | ns |
| Tahoe160 | 38 | 13.348 | -22.241 | 57.774 | 0.3223 | 0.3625 | ns |
| Tahoe183 | 45 | 45.545 | -1.393 | 58.214 | 0.0840 | 0.1512 | ns |
| Tahoe162 | 52 | 48.607 | 14.191 | 61.606 | 0.0098 | 0.0410 | * |
| Tahoe178 | 46 | 32.618 | -0.183 | 47.048 | 0.0645 | 0.1450 | ns |
| Tahoe163 | 51 | 41.291 | 10.524 | 67.235 | 0.0137 | 0.0410 | * |

**Table S18. Pairwise comparisons of relative abundances of amplicon sequence variants among frog body regions.** Following significant results of Kruskal-Wallis test (SV2 relative abundance: chi-squared = 35.697, df = 9, p < 0.001; SV1 relative abundance: chi-squared = 27.941, df = 9, p < 0.001), results of *post hoc* Dunn tests for significant differences in relative abundance of SV1 (family Rubritaleaceae) and SV2 (family Burkholderiaceae) in pairwise comparisons of frog body regions are shown; p-values were adjusted using Benjamini-Hochberg procedure to control the false discovery rate (FDR) with multiple comparisons; for significance, “***” = p ≤ 0.001, “**” = p ≤ 0.01, “*” = p ≤ 0.05, “ns” = not significant.

|  | **Comparison** | **Z** | **Unadjusted  P-value** | **Adjusted  P-value** | **Significance** |
| --- | --- | --- | --- | --- | --- |
| **SV2** | abdomen ↔ back | 3.406 | 0.000659 | 0.007419 | ** |
|  | abdomen ↔ cloaca | -0.054 | 0.956829 | 0.978575 | ns |
|  | back ↔ cloaca | -3.460 | 0.000540 | 0.008101 | ** |
|  | abdomen ↔ forefeet | 1.416 | 0.156630 | 0.293681 | ns |
|  | back ↔ forefeet | -1.989 | 0.046656 | 0.139967 | ns |
|  | cloaca ↔ forefeet | 1.471 | 0.141391 | 0.276635 | ns |
|  | abdomen ↔ hindfeet | -0.807 | 0.419382 | 0.555065 | ns |
|  | back ↔ hindfeet | -4.213 | 0.000025 | 0.001132 | ** |
|  | cloaca ↔ hindfeet | -0.753 | 0.451234 | 0.580158 | ns |
|  | forefeet ↔ hindfeet | -2.224 | 0.026149 | 0.090516 | ns |
|  | abdomen ↔ inner forelimbs | 0.875 | 0.381487 | 0.553771 | ns |
|  | back ↔ inner forelimbs | -2.531 | 0.011382 | 0.042682 | * |
|  | cloaca ↔ inner forelimbs | 0.929 | 0.352737 | 0.529105 | ns |
|  | forefeet ↔ inner forelimbs | -0.541 | 0.588276 | 0.735345 | ns |
|  | hindfeet ↔ inner forelimbs | 1.683 | 0.092442 | 0.218942 | ns |
|  | abdomen ↔ inner hindlimbs | 0.036 | 0.971211 | 0.971211 | ns |
|  | back ↔ inner hindlimbs | -3.370 | 0.000752 | 0.006770 | ** |
|  | cloaca ↔ inner hindlimbs | 0.090 | 0.928110 | 0.971278 | ns |
|  | forefeet ↔ inner hindlimbs | -1.380 | 0.167461 | 0.301430 | ns |
|  | hindfeet ↔ inner hindlimbs | 0.844 | 0.398903 | 0.560957 | ns |
|  | inner forelimbs ↔ inner hindlimbs | -0.839 | 0.401429 | 0.547403 | ns |
|  | abdomen ↔ outer hindlimbs | 0.370 | 0.711447 | 0.800378 | ns |
|  | back ↔ outer hindlimbs | -3.036 | 0.002397 | 0.011987 | * |
|  | cloaca ↔ outer hindlimbs | 0.424 | 0.671532 | 0.774844 | ns |
|  | forefeet ↔ outer hindlimbs | -1.047 | 0.295292 | 0.474576 | ns |
|  | hindfeet ↔ outer hindlimbs | 1.177 | 0.239033 | 0.413711 | ns |
|  | inner forelimbs ↔ outer hindlimbs | -0.505 | 0.613385 | 0.746009 | ns |
|  | inner hindlimbs ↔ outer hindlimbs | 0.334 | 0.738512 | 0.810562 | ns |
|  | abdomen ↔ snout | 3.081 | 0.002062 | 0.013258 | * |
|  | back ↔ snout | -0.325 | 0.745331 | 0.798569 | ns |
|  | cloaca ↔ snout | 3.135 | 0.001717 | 0.012879 | * |
|  | forefeet ↔ snout | 1.665 | 0.095991 | 0.215979 | ns |
|  | hindfeet ↔ snout | 3.889 | 0.000101 | 0.002269 | ** |
|  | inner forelimbs ↔ snout | 2.206 | 0.027388 | 0.088032 | ns |
|  | inner hindlimbs ↔ snout | 3.045 | 0.002327 | 0.013088 | * |
|  | outer hindlimbs ↔ snout | 2.711 | 0.006704 | 0.027426 | * |
|  | abdomen ↔ vocal sack | 1.917 | 0.055208 | 0.146140 | ns |
|  | back ↔ vocal sack | -1.489 | 0.136573 | 0.279354 | ns |
|  | cloaca ↔ vocal sack | 1.971 | 0.048682 | 0.136918 | ns |
|  | forefeet ↔ vocal sack | 0.501 | 0.616557 | 0.730133 | ns |
|  | hindfeet ↔ vocal sack | 2.725 | 0.006435 | 0.028960 | * |
|  | inner forelimbs ↔ vocal sack | 1.042 | 0.297378 | 0.461449 | ns |
|  | inner hindlimbs ↔ vocal sack | 1.881 | 0.059952 | 0.149881 | ns |
|  | outer hindlimbs ↔ vocal sack | 1.547 | 0.121786 | 0.260970 | ns |
|  | snout ↔ vocal sack | -1.164 | 0.244475 | 0.407459 | ns |
| **SV1** | abdomen ↔ back | -3.095 | 0.001970 | 0.017734 | * |
|  | abdomen ↔ cloaca | 0.469 | 0.638955 | 0.798694 | ns |
|  | back ↔ cloaca | 3.564 | 0.000366 | 0.008224 | ** |
|  | abdomen ↔ forefeet | 0.311 | 0.755596 | 0.829312 | ns |
|  | back ↔ forefeet | 3.406 | 0.000659 | 0.009891 | ** |
|  | cloaca ↔ forefeet | -0.158 | 0.874543 | 0.915220 | ns |
|  | abdomen ↔ hindfeet | 0.911 | 0.362162 | 0.626819 | ns |
|  | back ↔ hindfeet | 4.006 | 0.000062 | 0.002780 | ** |
|  | cloaca ↔ hindfeet | 0.442 | 0.658422 | 0.779710 | ns |
|  | forefeet ↔ hindfeet | 0.600 | 0.548517 | 0.747978 | ns |
|  | abdomen ↔ inner forelimbs | -0.442 | 0.658422 | 0.800783 | ns |
|  | back ↔ inner forelimbs | 2.653 | 0.007988 | 0.044935 | * |
|  | cloaca ↔ inner forelimbs | -0.911 | 0.362162 | 0.651891 | ns |
|  | forefeet ↔ inner forelimbs | -0.753 | 0.451232 | 0.700187 | ns |
|  | hindfeet ↔ inner forelimbs | -1.353 | 0.175945 | 0.416712 | ns |
|  | abdomen ↔ inner hindlimbs | 0.144 | 0.885219 | 0.905337 | ns |
|  | back ↔ inner hindlimbs | 3.239 | 0.001199 | 0.013494 | * |
|  | cloaca ↔ inner hindlimbs | -0.325 | 0.745330 | 0.838496 | ns |
|  | forefeet ↔ inner hindlimbs | -0.167 | 0.867439 | 0.929399 | ns |
|  | hindfeet ↔ inner hindlimbs | -0.767 | 0.443143 | 0.712194 | ns |
|  | inner forelimbs ↔ inner hindlimbs | 0.586 | 0.557573 | 0.737964 | ns |
|  | abdomen ↔ outer hindlimbs | -0.397 | 0.691381 | 0.797747 | ns |
|  | back ↔ outer hindlimbs | 2.698 | 0.006983 | 0.044888 | * |
|  | cloaca ↔ outer hindlimbs | -0.866 | 0.386412 | 0.644021 | ns |
|  | forefeet ↔ outer hindlimbs | -0.708 | 0.478789 | 0.718184 | ns |
|  | hindfeet ↔ outer hindlimbs | -1.308 | 0.190794 | 0.429286 | ns |
|  | inner forelimbs ↔ outer hindlimbs | 0.045 | 0.964018 | 0.964018 | ns |
|  | inner hindlimbs ↔ outer hindlimbs | -0.541 | 0.588274 | 0.756353 | ns |
|  | abdomen ↔ snout | -2.138 | 0.032493 | 0.121850 | ns |
|  | back ↔ snout | 0.956 | 0.338888 | 0.635415 | ns |
|  | cloaca ↔ snout | -2.607 | 0.009122 | 0.045610 | * |
|  | forefeet ↔ snout | -2.450 | 0.014303 | 0.064364 | ns |
|  | hindfeet ↔ snout | -3.050 | 0.002292 | 0.017189 | * |
|  | inner forelimbs ↔ snout | -1.696 | 0.089849 | 0.252700 | ns |
|  | inner hindlimbs ↔ snout | -2.283 | 0.022451 | 0.091846 | ns |
|  | outer hindlimbs ↔ snout | -1.741 | 0.081630 | 0.244890 | ns |
|  | abdomen ↔ vocal sack | -1.042 | 0.297376 | 0.581823 | ns |
|  | back ↔ vocal sack | 2.053 | 0.040114 | 0.138855 | ns |
|  | cloaca ↔ vocal sack | -1.511 | 0.130728 | 0.346045 | ns |
|  | forefeet ↔ vocal sack | -1.353 | 0.175945 | 0.439862 | ns |
|  | hindfeet ↔ vocal sack | -1.953 | 0.050780 | 0.163223 | ns |
|  | inner forelimbs ↔ vocal sack | -0.600 | 0.548517 | 0.771352 | ns |
|  | inner hindlimbs ↔ vocal sack | -1.186 | 0.235451 | 0.504538 | ns |
|  | outer hindlimbs ↔ vocal sack | -0.645 | 0.518866 | 0.753192 | ns |
|  | snout ↔ vocal sack | 1.096 | 0.272987 | 0.558382 | ns |

**Table S19. Comparison of SV1 and SV2 relative abundance within each frog body region.** Results of Wilcoxon signed rank tests for significant differences between relative abundances of SV1 (family Rubritaleaceae) and SV2 (family Burkholderiaceae) by frog body region are shown; p-values were adjusted using the Benjamini-Hochberg procedure to control the false discovery rate (FDR) with multiple comparisons; for significance, “***” = p ≤ 0.001, “**” = p ≤ 0.01, “*” = p ≤ 0.05, “ns” = not significant.

| **SampleType** | **Vstats** | **Estimate** | **Lower 95% CI** | **Upper 95% CI** | **P-value** | **Adjusted P-value** | **Significance** |
| --- | --- | --- | --- | --- | --- | --- | --- |
| back | 0 | -52.365 | -67.638 | -29.941 | 0.0039 | 0.0391 | * |
| snout | 1 | -33.975 | -59.223 | -9.974 | 0.0078 | 0.0391 | * |
| vocal sack | 18 | -5.079 | -43.766 | 31.060 | 0.6523 | 0.6523 | ns |
| inner forelimbs | 30 | 9.259 | -29.886 | 48.588 | 0.4258 | 0.4731 | ns |
| outer hindlimbs | 37 | 21.287 | -5.464 | 42.263 | 0.0977 | 0.1953 | ns |
| abdomen | 33 | 32.050 | -22.351 | 71.489 | 0.2500 | 0.3125 | ns |
| inner hindlimbs | 36 | 38.357 | -4.584 | 56.619 | 0.1289 | 0.2148 | ns |
| forefeet | 33 | 15.823 | -16.978 | 43.014 | 0.2500 | 0.3125 | ns |
| cloaca | 42 | 32.435 | 5.684 | 55.501 | 0.0195 | 0.0651 | ns |
| hindfeet | 39 | 36.652 | -2.237 | 71.452 | 0.0547 | 0.1367 | ns |

**Table S20. Differentially abundant amplicon sequence variants between ventral body regions and the dorsal back region.** Differentially abundant amplicon sequence variants identified by DESeq2 (*i.e.,* those that had significant log_2_ fold difference with p ≤ 0.05) are shown for each contrast; contrasts examined include abdomen – back, inner hindlimbs – back, forefeet – back, and hindfeet – back; “ASV” indicates the differentially abundant amplicon sequence variant and its associated taxonomy; the "base Mean'' is the mean DESeq2 normalized counts for the amplicon sequence variant across all samples; the "Estimate" is the log_2_ fold difference estimate for the specified contrast; the "Statistic" is the Wald statistic for the specified contrast; p-values are adjusted for multiple comparisons using the Benjamini-Hochberg procedure; “Signif.” indicates significance of adjusted p-values: “***” = p ≤ 0.001, “**” = p ≤ 0.01, “*” = p ≤ 0.05, “ns” = not significant.

| **Contrast** | **ASV** | **base Mean** | **Estimate** | **SE** | **Statistic** | **P-value** | **Adjust.  P-value** | **Signif.** |
| --- | --- | --- | --- | --- | --- | --- | --- | --- |
| abdomen ↔ back | SV56 F. Burkholderiaceae | 10.237 | 24.127 | 6.488 | 3.719 | 0.0002 | 0.0498 | * |
